# Endothelial pannexin 1 channels control inflammation by regulating intracellular calcium

**DOI:** 10.1101/750323

**Authors:** Yang Yang, Leon Delalio, Angela K Best, Edgar Macal, Jenna Milstein, Iona Donnelly, Ashley M. Miller, Martin McBride, Xiaohong Shu, Michael Koval, Brant E. Isakson, Scott R. Johnstone

**Affiliations:** Robert M. Berne Cardiovascular Research Center, University of Virginia School of Medicine; Department of Pharmacology, Dalian Medical University, Dalian 116044, China; British Heart Foundation Cardiovascular Research Centre, College of Medical, Veterinary and Life Sciences, University of Glasgow, Glasgow G12 8TA, UK; Division of Pulmonary, Allergy, Critical Care and Sleep Medicine, Department of Medicine, Emory University School of Medicine, Atlanta, GA 30322, USA; Department of Cell Biology, Emory University School of Medicine, Atlanta, GA 30322, USA; Department of Molecular Physiology and Biophysics, University of Virginia School of Medicine

**Keywords:** Pannexin, Panx1, interleukin-1β, NFκβ, ATP, calcium, gene regulation, channel, inflammation, endothelial cell, smooth muscle cell

## Abstract

**In Brief:** Interleukine-1 beta (IL-1β) has been identified as a critical factor that contributes to the inflammatory response in cardiovascular disease (e.g., atherosclerosis). Pannexin 1 (Panx1) channel activity in endothelial cells regulates localized inflammatory cell recruitment. In response to prolonged tumor necrosis factor alpha (TNF) treatment, Yang et al. found that the Panx1 channel is targeted to the plasma membrane, where it facilitates an increase in intracellular calcium to control the production and release of cytokines including IL-1β.

**GRAPHICAL ABSTRACT:** 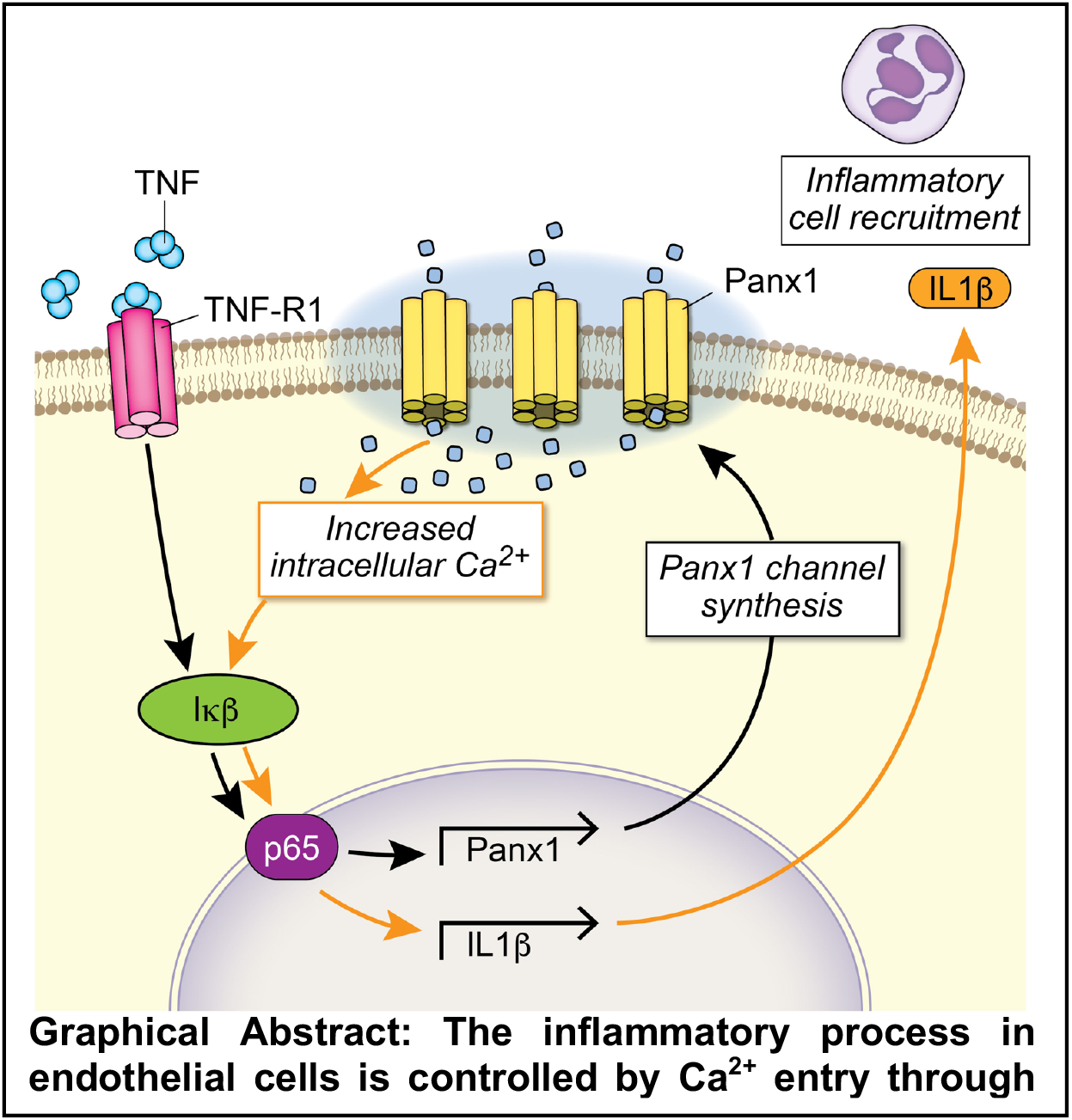

**Abstract:** The proinflammatory cytokine IL-1β is a significant risk factor in cardiovascular disease that can be targeted to reduce major cardiovascular events. IL-1β expression and release are tightly controlled by changes in intracellular Ca^2+^. In addition, purinergic signaling through ATP release has also been reported to promote IL-1β production. Despite this, the mechanisms that control IL-1β synthesis and expression have not been identified. The pannexin 1 (Panx1) channel has canonically been implicated in ATP release, especially during inflammation. However, resolution of purinergic signaling occurs quickly due to blood flow and the presence of ectonucleotidases. We examined Panx1 in human endothelial cells following treatment with the pro-inflammatory cytokine tumor necrosis alpha (TNF). In response to long-term TNF treatment, we identified a dramatic increase in Panx1 protein expression at the plasma membrane. Analysis by whole transcriptome sequencing (RNA-seq), qPCR, and treatment with specific kinase inhibitors, revealed that TNF signaling induced NFκβ-associated Panx1 transcription. Genetic inhibition of Panx1 reduced the expression and secretion of IL-1β. We initially hypothesized that increased Panx1-mediated ATP release acted in a paracrine fashion to control cytokine expression. However, our data demonstrate that IL1-β expression was not altered after direct ATP stimulation, following degradation of ATP by apyrase, or after pharmacological blockade of P2 receptors. These data suggest that non-purinergic pathways, involving Panx1, control IL-1β production. Because Panx1 forms a large pore channel, we hypothesized it may act to passively diffuse Ca^2+^ into the cell upon opening to regulate IL-1β. High-throughput flow cytometric analysis demonstrated that TNF treatments lead to elevated intracellular Ca^2+^. Genetic or pharmacological inhibition of Panx1 reduced TNF-associated increases in intracellular Ca^2+^, and IL-1β transcription. Furthermore, we found that the Ca^2+^-sensitive NFκβ-p65 protein failed to localize to the nucleus after genetic or pharmacological block of Panx1. Taken together, our study provides the first evidence that intracellular Ca^2+^ regulation via the Panx1 channel induces a feed-forward effect on NFκβ to regulate IL-1β synthesis and release in endothelium during inflammation.

## INTRODUCTION

Sustained inflammatory responses critically regulate the pathogenesis of endothelial dysfunction and atherosclerosis (Libby and Hansson, 2015; Libby et al., 2013). The release of tumor necrosis factor alpha (TNF) enhances inflammation in atherosclerosis and TNF concentrations are associated with elevated risk of atherothrombosis and resulting major adverse cardiovascular events (Barath et al., 1990; Ridker, 2013; Ridker et al., 2000). While many studies have focused on the effect of TNF on inflammatory cells (Bradley, 2008), TNF has also been shown to induce production of pro-inflammatory cytokines in endothelial cells (ECs) (Imaizumi et al., 2000; Perrot-Applanat et al., 2011). In the presence of TNF, ECs synthesize and release pro-inflammatory cytokines and chemokines that enhance the inflammatory response, correlating with a high risk of vascular injury (Perrot-Applanat et al., 2011; Viemann et al., 2006). Targeting inflammation, e.g. using TNF antagonists, leads to reductions in cytokine expression by ECs and can reduce atherosclerotic lesion formation (Alexander et al., 2012; Di Minno et al., 2011; Gabay et al., 2016; Virone et al., 2019). Thus, defining critical EC signaling pathways may help identify therapeutic targets. Among these potential targets, interleukin-1β (IL-1β) is widely considered to be a highly active and essential regulator of the pathogenesis of human atherosclerotic disease progression and susceptibility to atherothrombosis (Ridker et al., 2012). Therapeutically targeting IL-1β decreases its activity and is associated with a reduced expression of multiple pro-inflammatory cytokines, including interleukin-6 (IL-6), which have been implicated as a potential causal pathway for atherosclerotic events (Ridker et al., 2017; Solomon et al., 2018). In the recent Canakinumab Anti-inflammatory Thrombosis Outcome Study (CANTOS), IL-1β neutralization by canakinumab reduced inflammation and reduced major adverse cardiovascular events associated with atherothrombosis in high-risk patients (Ridker et al., 2017; Ridker et al., 2012). While other pro-inflammatory cytokines such as IL-6 have been implicated in atherosclerosis, clinical trials targeting this did not result in marked improvements in patient risk, highlighting the importance of targeting specific pro-inflammatory markers (Ridker et al., 2019; Ridker et al., 2018). The efficacy of specific IL-1β blockade in inflammation highlights the need to elucidate the molecular mechanisms that regulate its synthesis and release.

Adenosine triphosphate (ATP), is increasingly recognized as an important factor in the regulation of the inflammatory process, leading to activation of the inflammasome (Maitre et al., 2015; Mariathasan et al., 2006; Qu et al., 2011), and multiple studies have demonstrated its association with increases in IL-1β synthesis and release (Kanjanamekanant et al., 2013; Mehta et al., 2001). While some studies suggest that ATP alone is capable of increasing IL-1β synthesis and release (Kanjanamekanant et al., 2013), a “two-signal” model of production followed by later activation has emerged for IL-1β (Cullen et al., 2015; Mehta et al., 2001). In this model, ATP acts to enhance the magnitude and velocity of posttranslational processing of pro-IL-1β, yielding the bioactive molecule, which is released into the extracellular space (Cullen et al., 2015). However, in this model, IL-1β synthesis is not controlled via ATP, rather through pathways stimulated via pathogen-associated molecules such as LPS (Cullen et al., 2015; Stoffels et al., 2015). Release of ATP from cells can also signal locally through paracrine receptors (e.g. P2X7) to promote K^+^ release leading to uptake of extracellular Ca^2+^, which is associated with an increase in the expression and release of cytokines including IL-1β (Jantaratnotai et al., 2009; Wilson et al., 2007; Yaron et al., 2015). Further data suggests that chelation of intracellular Ca^2+^ inhibits the processing and release of IL-1β, suggesting that an influx of extracellular Ca^2+^ is also centrally linked to IL-1β production (Ainscough et al., 2015; Brough et al., 2003). Despite this, the source and regulation of increased intracellular Ca^2+^ has not been rigorously defined (Ainscough et al., 2015; Brough et al., 2003).

Pannexin 1 (Panx1) forms large, non-selective plasma membrane channels that permit the movement of molecules and ions, including ATP to the extracellular space and Ca^2+^ from ER (Chiu et al., 2018; Vanden Abeele et al., 2006). Panx1 channels at the plasma membrane can facilitate multiple physiological and pathophysiological processes including vascular constriction, apoptosis, tumor cell metastasis, and neuronal communication (Billaud et al., 2011; Chekeni et al., 2010; Orellana et al., 2011; Thompson et al., 2008). Recently, we identified that endothelial Panx1 channel opening and ATP release promotes leukocyte recruitment (Lohman et al., 2015) that plays a fundamental role in inflammation and tissue damage within ischemic stroke (Good et al., 2018b). There is increasing evidence that blocking Panx1 channels may control inflammasome activation, inflammatory cytokine release and inflammatory cell recruitment (Albalawi et al., 2017; Lappas, 2014; Pelegrin, 2008). However, the mechanisms underlying this response have not been described. This led us to hypothesize that Panx1 signaling may be involved in the control of IL-1β production and secretion by ECs. We report here that EC Panx1 is a direct target of the TNF signaling pathway and demonstrate for the first time that Panx1 channels facilitate the transport of extracellular Ca^2+^ to promote a feed forward effect on the synthesis of IL-1β.

## RESULTS

### TNF induces Panx1 expression and membrane targeting of Panx1 channels in endothelial cells

In human umbilical vein endothelial cells (HUVECs), we initially demonstrate that prolonged exposure to TNF (2.5ng/mL) for 5 hr and 24 hr leads to an increase in the transcription of Panx1, measured by qRT-PCR (**Figure 1A**). Increases in Panx1 mRNA levels correlate with significant increases in protein expression of Panx1 at 5 hr, measured by immunoblotting (**Figure 1B**). The multiple banding pattern for the Panx1 protein observed in immunoblots represents differential Panx1-glycosylation (Panx1-Gly) isoforms, referred to as Panx1-Gly0, -Gly1 and -Gly2 (Boassa et al., 2007; Penuela et al., 2007; Penuela et al., 2008). Interestingly, we observed that, at 5 hr, TNF treatments primarily increase only Panx1-Gly0 and Panx1-Gly1 isoforms, which were ablated by co-treatment with the protein synthesis inhibitor cycloheximide (CHX, **Figure 1B** and **Supplemental Figure 1A**), suggesting that these isoforms represent newly synthesized Panx1. To determine the specificity of TNF to Panx1 expression, we investigated the Cx43 gap junction protein in HUVECs, which did not increase following TNF stimulation and was inhibited following CHX treatments (**Figure 1B**). Following 24 hr TNF treatment, we identified significant upregulation of the Panx1-Gly2 isoform (**Supplemental Figure 1B**). Increasing the concentration of TNF treatments did not enhance Panx1 expression from 2.5 ng/mL at either 5 hr or 24 hr and did not alter expression of Cx43 (**Supplemental Figure 1A-C**). In addition, Panx1-Gly2 isoforms were not reduced by CHX (**Figure 1B**), suggesting that these represent a more stable isoform of Panx1. To investigate this, HUVECs were treated with TNF for 24 hr, washed in fresh media without TNF, and cultured for a further 8 days. Analysis by immunoblot demonstrates that while the Panx1-Gly0/1 isoforms are lost within 24 hr of removal of TNF, the Panx1-Gly2 isoforms remain significantly increased from control for 24 hr after removal of TNF and remain elevated for a further 5 days (**Figure 1C**). This demonstrates a stable, mature form of Panx1 that has a significant residence time at the plasma membrane, making it more stable than connexin proteins.

**Figure 1.**
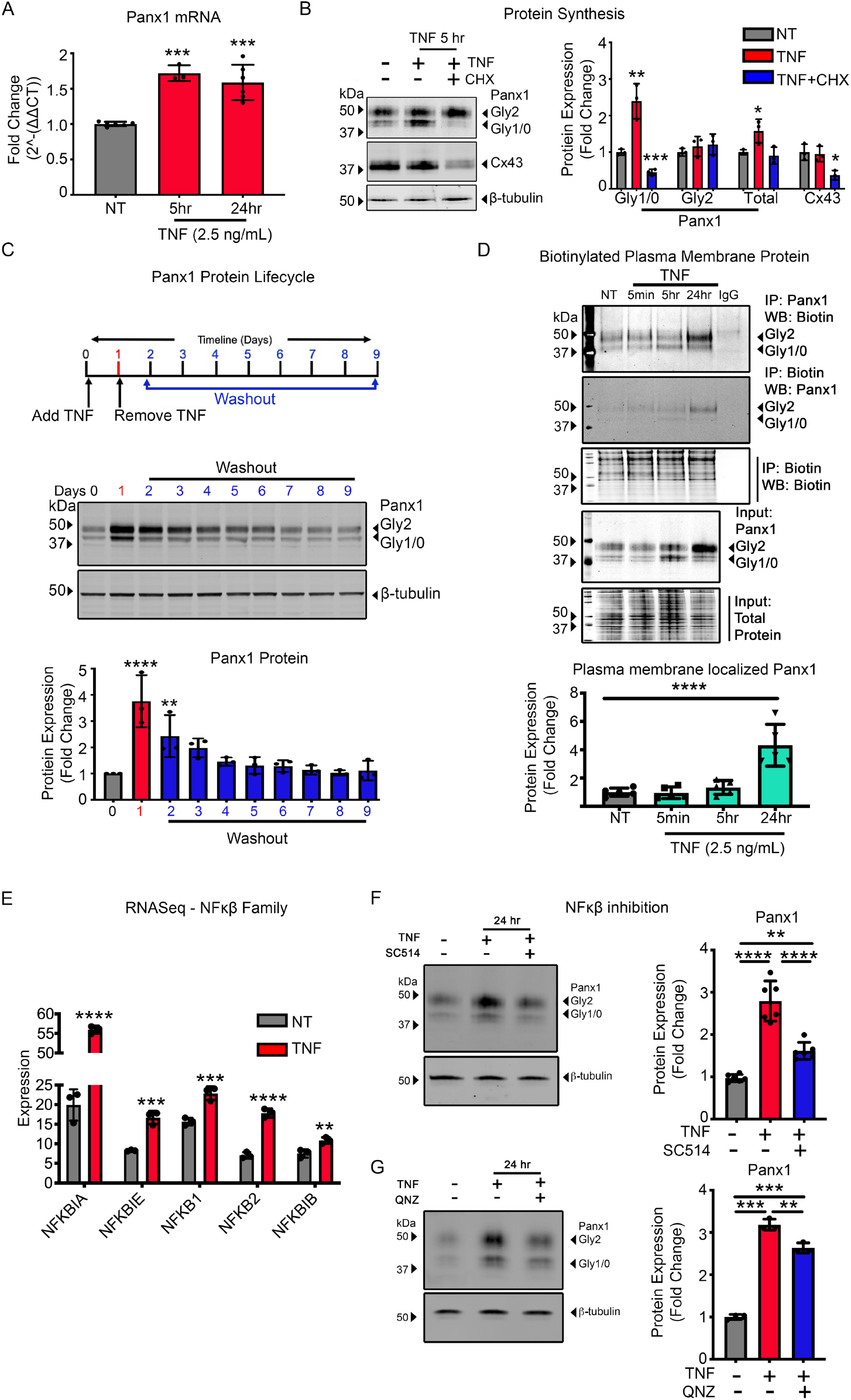
TNF transcriptional regulation of Panx1 through the NFκβ pathway promotes protein synthesis and plasma membrane trafficking. (A) Taqman qRT-PCR of RNA extracted from HUVECs treated with TNF (2.5 ng/mL) for 5 and 24 hr. The mean of Panx1 expression was normalized to control and calculated to 2^-ΔΔCT. Each error bar was performed in triplicates in addition to the technical triplicates. (B) Representative immunoblots of pannexin 1 (Panx1) and Connexin 43 (Cx43) in HUVECs pretreated with cycloheximide (CBX) 30 minutes and subsequent TNF (2.5 ng/mL) treatment for 24 hr. Quantification of Panx1 expression was normalized to β-tubulin (n=3). (C) Upper panel schematic of HUVECs treatment approach to assess Panx1 protein lifecycle. Black arrowhead marks the time TNF was added and removed. The blue scale marks cells maintained in 0.1%-M200 (no TNF) collected each 24 hr after TNF removal. Lower panel, representative immunoblots of Panx1 and β-tubulin expression and quantification of Panx1 expression under these treatment conditions (n=3). (D) Representative immunoblots of HUVECs treated with TNF (2.5 ng/mL) for time course 5 minutes, 5 and 24 hr, and subsequent immunoprecipitation of cell surface biotinylated membrane proteins. Plasma membrane localized Panx1 expression was normalized to biotin-labeled total protein (n=5). (E) RNA sequence performed on HUVECs treated with TNF (2.5 ng/mL) for 24 hr. The expression of five genes in NFκβ family are shown with each bar represents mean±SD for triplicates (n=3). (F) Represent immunoblots of TNF (2.5 ng/mL) induced HUVECs in presence or absence of inhibitors: inhibitor of nuclear factor kappa-B kinase-2 (IKK2) 100 μM SC514 (n=5) or NFκβ inhibitor 10 μM QNZ (n=3) for 24 hr. Statistical analyses were performed by one-way or two-way ANOVA with either Dunnett or Tukey’s multiple comparison test, *P<0.05, **P<0.01, ***P<0.001, ****P<0.001.

Previous studies have suggested that Panx1-Gly2 isoforms represent the mature Panx1 channels within the plasma membrane (Boassa et al., 2007). We therefore used a membrane protein biotinylation pull-down approach to isolate plasma membrane proteins and identified that Panx1-Gly2 isoforms are primarily localized within the plasma membrane 24 hr after TNF treatment (**Figure 1D**). We further demonstrate that Panx1 expression by cultured human vascular smooth muscle cells (SMC) is unaltered by TNF treatment (**Supplemental Figure 1D**). Previous studies have found TNF to reduce cell viability (Zhou et al., 2017). However, 24 hr TNF treatments ranging from 2.5-100 ng/mL for 24 hr did not alter HUVEC viability as measured by intracellular ATP levels, caspase-3 cleavage, or by cell morphology (**Supplemental Figure 1E-G**).

TNF signaling is associated with activation of NFκβ-mediated gene regulation. Whole-transcriptome RNA sequencing (RNA-seq) experiments confirmed that TNF treatment of HUVEC significantly upregulated NFκβ genes NFKBIA, NFKBIE, NFKB1, NFKB2, NFKB1B (**Figure 1E**). To define a role for NFκβ-activated pathways in TNF-induced increases in Panx1 expression, HUVECs were pre-treated with NFκβ-inhibitors SC514 and QNZ. Both SC514 and QNZ, significantly ablated TNF-induced increases in Panx1 expression at 24 hr (**Figure 1F-G**). RNA-seq results demonstrate that TNF (2.5 ng/mL) does not alter mitogen activated protein kinase (MAPK) transcription, and MAPK inhibition failed to reduce TNF-associated Panx1 upregulation (**Supplemental Figure 1H-I**). This suggests that TNF specifically stimulates a NFκβ-mediated upregulation of Panx1 proteins, leading to plasma membrane localization of Panx1 channels in ECs.

### Increased Panx1 membrane targeting is associated with the transcription and release of specific pro-inflammatory cytokines

Panx1 has been associated, but not directly implicated, in the release of molecules associated with inflammation (Chen et al., 2019; Lohman et al., 2015; Sharma et al., 2018). To investigate whether the increased Panx1 at the plasma membrane is directly associated with cytokine production and release, we first used genetic inhibition of Panx1. Panx1-siRNA reduced the basal expression of Panx1 in HUVECs and inhibited increases in Panx1 in response to TNF (**Figure 2A-B**). Cytokine arrays used to assay the media from HUVEC cells showed that TNF treatment altered the expression of a number of cytokines (**Supplemental Figure 2**). We selected key pro-inflammatory cytokines associated with atherosclerosis such as IL-1β and CXCL10, IL8 and MCP-1 which were found to be increased in response to TNF treatment in HUVEC, unlike MIF and Basigin which were not markedly increased by TNF (**Figure 2C**). siRNA knockdown of Panx1 in HUVECs was associated with a reduction in the release of IL-1β and CXCL10 (**Figure 2C, Supplemental Figure 2**). Notably, Panx1 siRNA treated cells displayed a significantly reduced transcription of IL-1β and CXCL10 but showed no differences in IL8 and CCL2 (**Figure 2D-E**). Panx1 siRNA and TNF-treatments also had no effect on MIF or CD147 (**Figure 2F**). These data suggest that Panx1 critically regulates the control of specific inflammatory cytokines including IL-1β and CXCL10. While multiple other cytokines and chemokines have been found to be increased in atherosclerosis, clinical trials targeting these pathways by treatments including low dose methotrexate has not proven to be effective in reducing the burden of disease in patients (Ridker et al., 2019). Based on this, we focused on mechanisms controlling IL-1β, since directly targeting IL-1β is linked with a reduction in patient risk (Ridker et al., 2017). Critically, understanding the molecular mechanisms controlling IL-1β expression may provide avenues for future therapeutic intervention (Brown et al., 2015; Dinarello, 2010; Libby and Hansson, 2015; Ridker, 2013, 2019).

**Figure 2.**
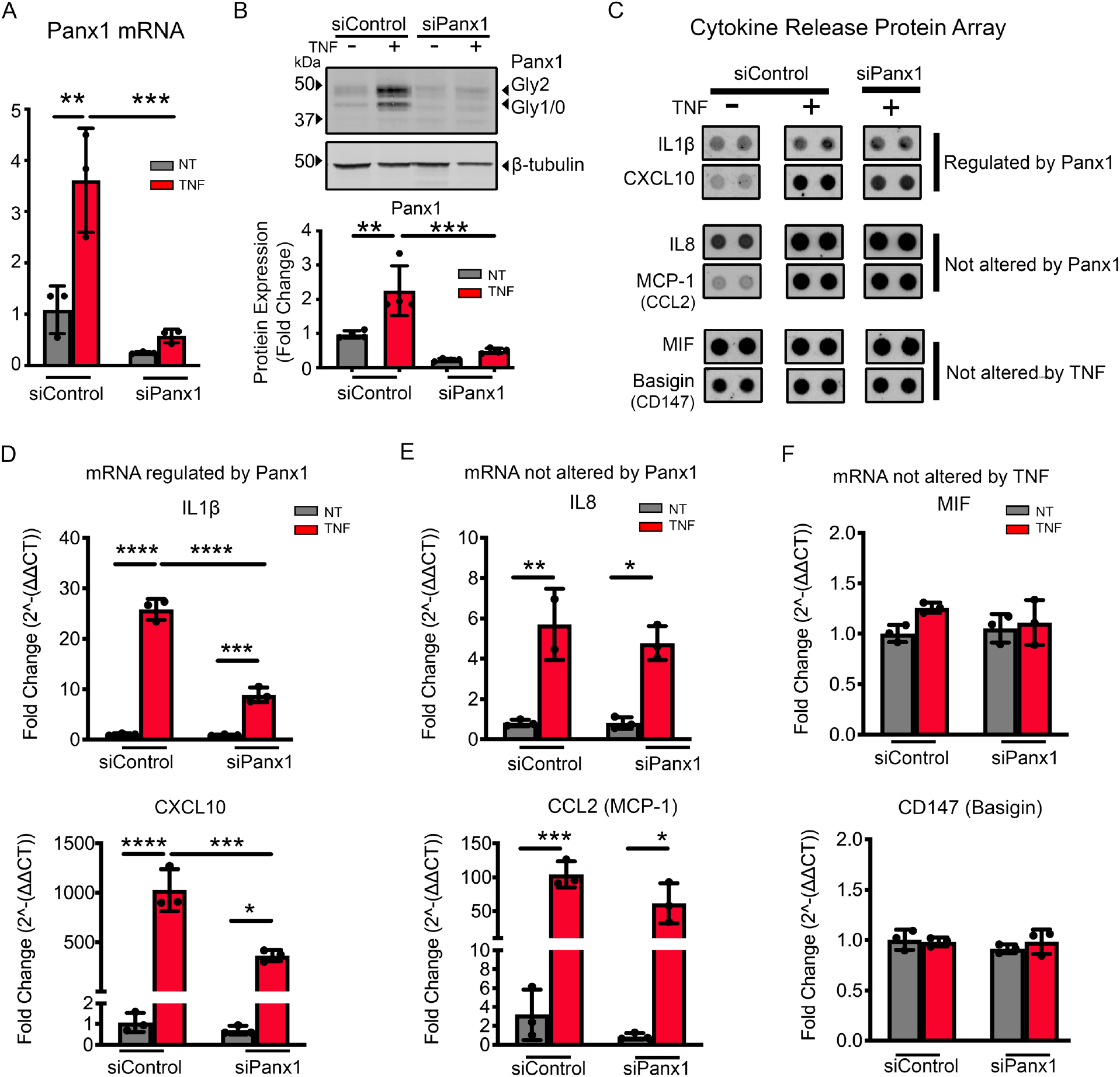
Panx1 controls transcription of selective inflammatory cytokines. (A) qRT-PCR analysis of Panx1 in HUVECs transfected with control siRNA (siControl) or siRNA Panx1 for 48 hr followed with 24 hr TNF (2.5 ng/mL) treatment. Each reaction was performed in triplicates in addition to the technical triplicates (n=3). (B) Representative immunoblot of Panx1 in HUVECs transfected with siControl or siRNA Panx1 for 48 hr followed 24 hr TNF (2.5 ng/mL) treatment. Panx1 expression was normalized to β-tubulin and expressed as fold change (n=4). (C) Cell media collected from HUVECs transfected with siControl or siRNA Panx1 for 48 hr followed with 24 hr TNF (2.5 ng/mL) treatment were incubated with human XL cytokine array (n=2). Representative cytokine spot duplicates for each group were shown with the relative mean measurement. qRT-PCR analysis the expression of IL-1β and CXCL10 (**D**), IL8 and CCL2 (**E**), and MIF and CD147 (**F**) in HUVECs transfected with control siRNA (siControl) or siRNA Panx1 for 48 hr followed with 24 hr TNF (2.5 ng/mL) treatment. Cytokines were normalized to control and calculated to 2^-ΔΔCT, then expressed as fold change. Each group was performed in triplicates in addition to the technical triplicates (n=3). Statistical analyses were performed by one-way or two-way ANOVA with either Dunnett or Tukey’s multiple comparison test, *P<0.05, **P<0.01, ***P<0.001, ****P<0.001.

### Purinergic signaling is not a key regulator of TNF-induced IL-1β in endothelial cells

Based on the known role for Panx1 and ATP release, which has been linked with IL-1β regulation, we investigated ATP release from HUVECs following treatment with TNF and the corresponding effects of ATP on IL-1β. ATP release from HUVECs was measured in media following TNF treatment at doses ranging from 2.5-100 ng/mL. All concentrations of TNF produced similar increases in ATP release from HUVECs (**Figure 3A**), which was significantly reduced following genetic inhibition of Panx1 (**Figure 3B**). While previous studies have correlated ATP treatment with regulation of IL-1β, this has typically required ATP analogues to be applied at concentrations ranging from 100 μM – 5 mM (Cullen et al., 2015; Kanjanamekanant et al., 2013; Mehta et al., 2001). Our data demonstrate that the HUVECs release only 10-20 nM ATP once exposed to TNF (2.5 ng/mL) for 24 hr (**Figure 3C**). At these concentrations (10 nM - 100 μM), the ATP analogue ATPγS failed to increase IL-1β transcription (**Figure 3D**). However, we did observe a small increase in transcription of IL-1β at 100 μM ATPγS, which could be ablated by pre-treating cells with the P2X inhibitor suramin (**Figure 3E**). Further data demonstrate that promoting the degradation of released ATP using apyrase or inhibition of the P2 receptors using suramin, did not significantly alter the expression of either Panx1 or IL-1β in the presence of TNF (**Figure 3F-G**). Thus, the NFκβ-induced increase in Panx1 expression at the plasma membrane, while crucial for IL-1β cytokine production, was not regulated by an autocrine ATP release from the EC.

**Figure 3.**
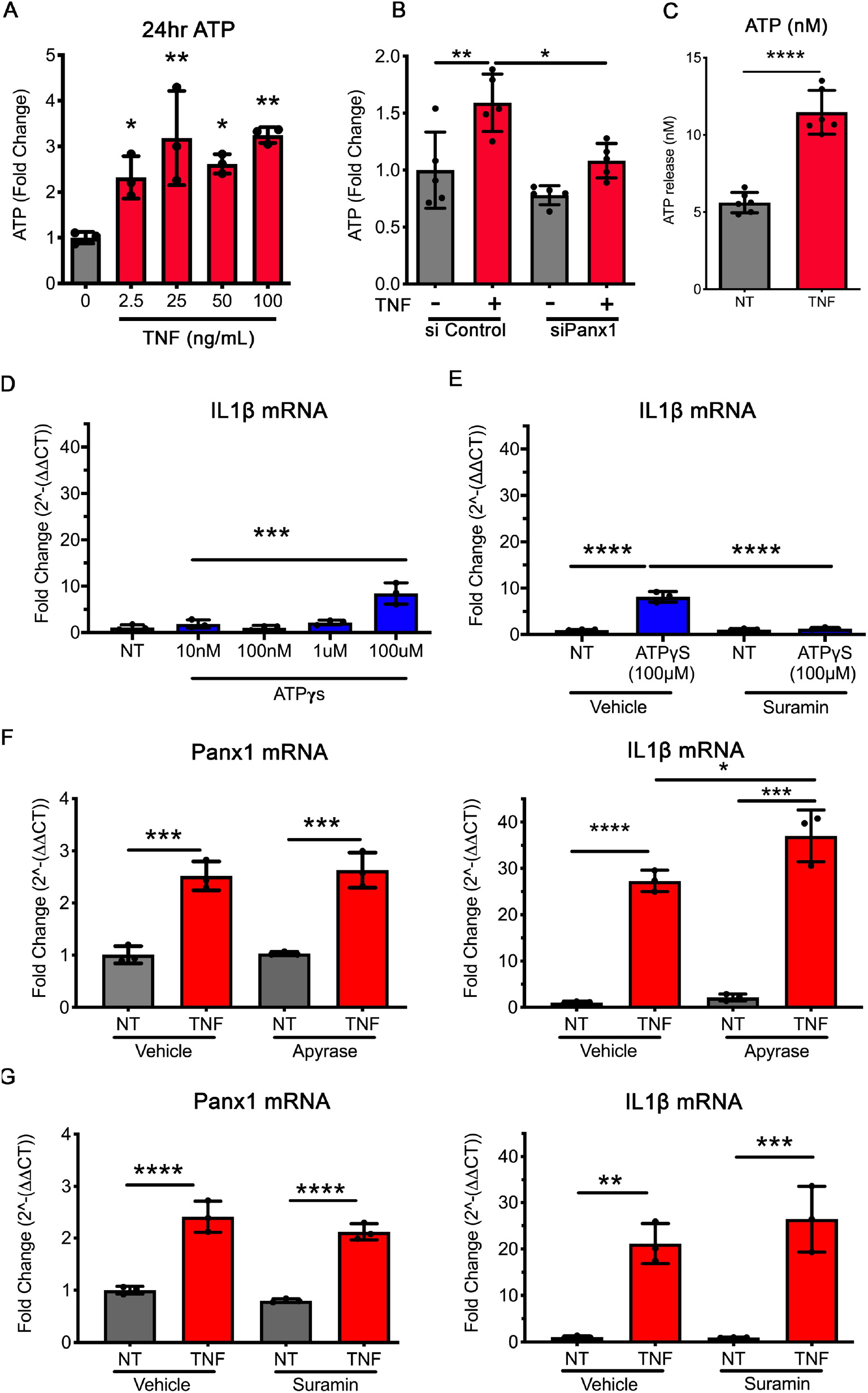
ATP release through Panx1 channel opening is not associated with IL-1β regulation. (A) ATP release measured and quantified as fold change from control (0 ng/mL TNF) to HUVECs pretreated with TNF (0, 2.5, 25, 50, 100 ng/mL) for 24 hr then stimulated with TNF (10 ng/mL) stimulated for 5 minutes (n=3). (B) ATP release measured and quantified as fold change from control (siControl or siPanx1, 0 ng/mL TNF) with 24 hr incubation of TNF (2.5 ng/ml) in response to 5 minutes TNF (10 ng/mL) stimulation (n=5). (C) Calculated mean extracellular ATP concentration from 24 hr TNF pretreatment HUVECs in response to TNF (10 ng/mL) for 5 minutes (n=6). (D) qRT-PCR analysis of IL-1β expression in HUVECs applied to a dose response for exogenous ATPγs (10, 100 nM, 1 and 100 μM), a stabilized ATP substrate to assess the potential effect on IL-1β upregulation (n=3). Data were normalized to control and calculated to 2^-ΔΔCT, then expressed as fold change. Each group was performed in triplicates in addition to the technical triplicates (n=3). (E) qRT-PCR analysis of IL-1β in HUVECs treated with ATPγs (100 μM), in the presence of suramin to (100 μM) to block P2X receptors (n=3). Data were normalized to control and calculated to 2^-ΔΔCT, then expressed as fold change. Each group was performed in triplicates in addition to the technical triplicates (n=3). (F) qRT-PCR analysis of Panx1 and IL-1β in HUVECs pretreated with Apyrase (1UN/mL), to degrade extracellular ATP, then with TNF (2.5 ng/mL) for 24 hr (n=3). Data were normalized to control and calculated to 2^-ΔΔCT, then expressed as fold change. Each group was performed in triplicates in addition to the technical triplicates (n=3). (G) qRT-PCR analysis of Panx1 and IL-1β in HUVECs pretreated with Suramin (100 μM), to block P2X receptor activity, then with TNF (2.5 ng/mL) for 24 hr (n=3). Data were normalized to control and calculated to 2^-ΔΔCT, then expressed as fold change. Each group was performed in triplicates in addition to the technical triplicates (n=3). Statistical analyses were performed by one-way or two-way ANOVA with either Dunnett or Tukey’s multiple comparison test, *P<0.05, **P<0.01, ***P<0.001, ****P<0.001.

### TNF increases in intracellular Ca^2+^ which is regulated by the functional Panx1 channels

Increased intracellular Ca^2+^ is associated with the control of IL-1β synthesis (Ainscough et al., 2015), and Panx1 channels have been suggested to allow the passage of Ca^2+^ from the extracellular environment into the cytosol, albeit this has never been demonstrated (Penuela et al., 2013; Vanden Abeele et al., 2006). To investigate whether Panx1 plays a role in the control of intracellular Ca^2+^ we used a high-throughput, flow cytometric approach (Vines et al., 2010), to measure intracellular Ca^2+^ in HUVECs. We found that intracellular calcium concentration ([Ca^2+^]_i_) was only increased after 24 hr TNF treatment, a time that correlates with the increases in Panx1 membrane localization (**Figure 4A**). To determine whether the increase in [Ca^2+^]_i_ was due to extracellular Ca^2+^ (e.g., through the increased number of Panx1 channels), Ca^2+^-free Krebs ringer was used. This experiment demonstrated a significant decrease in EC [Ca^2+^]_i_ (**Figure 4B**). We further demonstrated that increased intracellular Ca^2+^ is directly related to IL-1β production, since cells loaded with EGTA-AM in the media reduced [Ca^2+^]_i_ which corresponded to a loss of TNF-induced IL-1β transcription (**Figure 4C-D**). To more specifically assess whether there was a role for Panx1 in regulating EC [Ca^2+^]_i_, we used the Panx1-specific channel blocking peptide PxIL2P to pharmacologically inhibit Panx1 channels (Billaud et al., 2015; Good et al., 2018a). Pre-treatment of HUVECs with PxIL2P reduced [Ca^2+^]_i_ and IL-1β transcription in response to 24 hr TNF treatment (**Figure 4E-F**). Similar results were found using genetic inhibition of Panx1, which reduced TNF-associated increases in [Ca^2+^]_i_ (**Figure 4G)**and IL-1β (**Figure 2A**). In line with these observations, there was also a decrease in monocyte binding to EC with no change in THP1-Panx1 expression (**Supplemental Figure 3**). Lastly, levels of [Ca^2+^]_i_ and IL-1β mRNA were reduced 24 hr after the removal of TNF (**Figure 2H-I**), which correlated with the same decrease in Panx1 expression at the plasma membrane seen in **Figure 1C**. This data provides additional evidence that increased Panx1 channels in EC after TNF stimulation permit the passive diffusion of extracellular Ca^2+^ into the cell, possibly to regulate IL-1β production.

**Figure 4.**
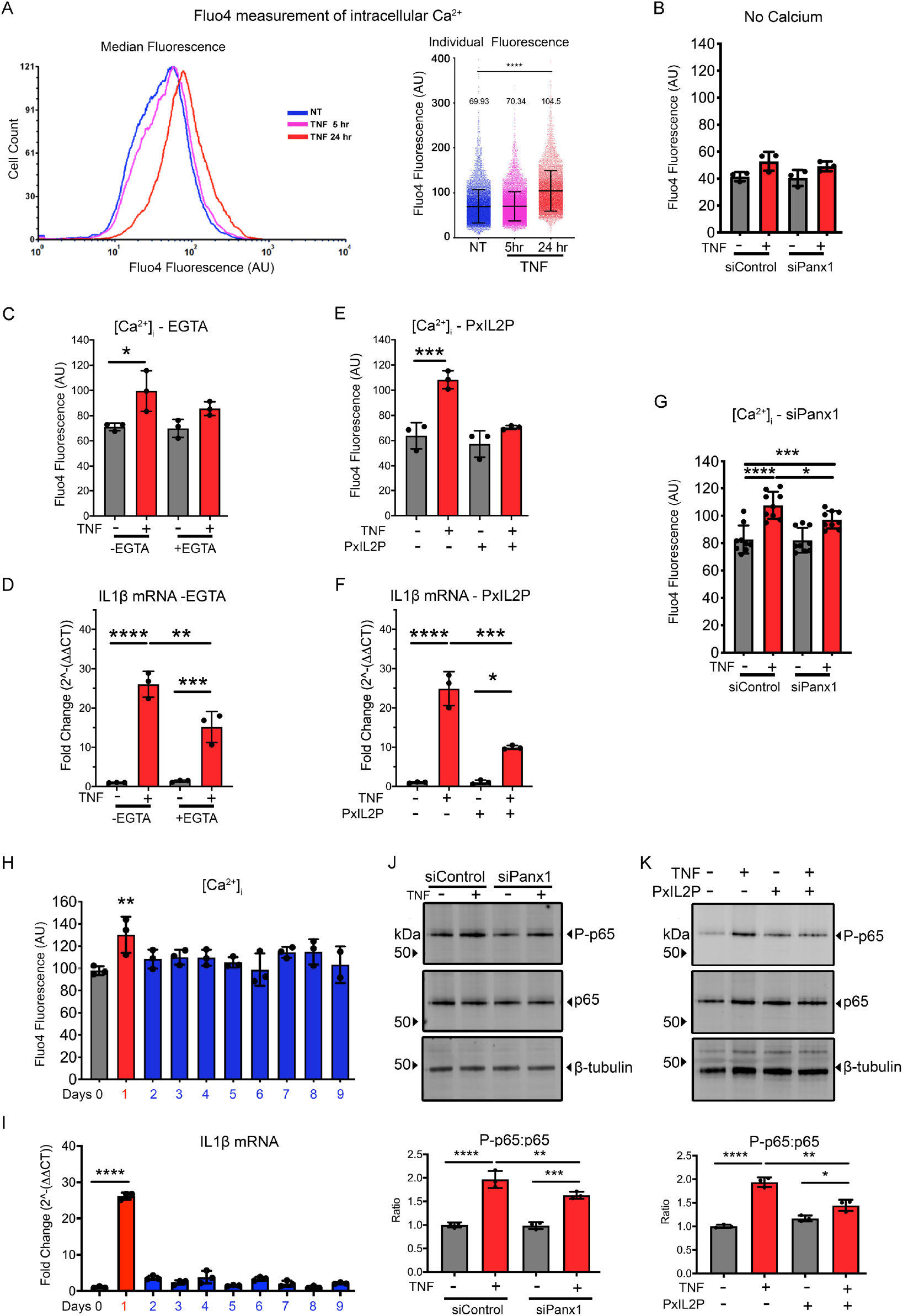
Panx1 facilitates increased intracellular Ca^2+^ associated with inflammatory regulation. (A) Represent [Ca^2+^]_i_ traces evoked in HUVECs treated with TNF (2.5 ng/mL) for 24 hr (red) compared to in control (blue) and 5 hr treatment (magenta). Cells were loaded with Fluo-4 AM (1 μM) and analyzed on BD FACScanto II flow cytometry. Graphs of median Fluo4 fluorescence of [Ca^2+^]_i_, were quantified for average individual fluorescence compared with control as indicated (n=3). (B) Flow cytometric measurement of median [Ca^2+^]_i_ in HUVEC in calcium free Krebs solution siRNA treatments (siControl, siPanx1) transfected cells treated with TNF (2.5 ng/mL) for 24 hr (n=3). (C) Flow cytometric measurement of median [Ca^2+^]_i_ in HUVEC treated with TNF (2.5 ng/mL) for 24 hr in presence of a chelator of calcium EGTA-AM (n=3). (D) qRT-PCR analysis of IL-1β expression in HUVECs treated with TNF (2.5 ng/mL) for 24 hr in the presence of a chelator of calcium EGTA-AM (n=3). Data were normalized to control and calculated to 2^-ΔΔCT, then expressed as fold change. Each group was performed in triplicates in addition to the technical triplicates (n=3). (E) Flow cytometric measurement of median [Ca^2+^]_i_ in HUVEC pretreated with PxIL2P peptide followed by TNF (2.5 ng/mL) treatment for 24 hr (n=3). (F) qRT-PCR analysis of IL-1β expression in HUVECs pretreated with Panx1 IL2 peptide followed by TNF (2.5 ng/mL) treatment for 24 hr. Data were normalized to control and calculated to 2^-ΔΔCT, then expressed as fold change. Each group was performed in triplicates in addition to the technical triplicates (n=3). (G) Flow cytometric measurement of median [Ca^2+^]_i_ in HUVECs transfected with siPanx1 followed by TNF (2.5 ng/mL) stimulation (n=3). (H) Flow cytometric measurement of median [Ca^2+^]_i_ in HUVECs treated with TNF for 24 hr as per experimental set up illustrated in Figure 1C schematic. (I) qRT-PCR analysis of IL-1β expression in HUVECs treated with TNF for 24 hr, then washed out for 8 days, as per experimental set up illustrated in Figure 1C schematic. Data were normalized to control and calculated to 2^-ΔΔCT, then expressed as fold change. Each group was performed in triplicates in addition to the technical triplicates (n=3). (J) Represent immunoblots of HUVECs treated with TNF (2.5 ng/mL) for 24 hr post-transfected with siRNA (siControl or siPanx1) for 48 hr. β-tubulin was used as a loading control. The relative changes of P-p65 and p65 expression were calculated in comparison with siControl no treatment (n=3). (K) Represent immunoblots of HUVECs pretreated with PxIL2P peptide followed with TNF (2.5 ng/mL) stimulation for 24 hr. β-tubulin was used as a loading control. The relative changes of P-65 and p65 expression were calculated in comparison with control no treatment (n=3). Statistical analyses were performed by one-way or two-way ANOVA with either Dunnett or Tukey’s multiple comparison test, *P<0.05, **P<0.01, ***P<0.001, ****P<0.001.

Because increased intracellular Ca^2+^ have been reported to result in phosphorylation of the NFκβ protein p65 (P-p65), increasing its transcriptional activity (Martin et al., 2001), we aimed to assess the role of Panx1 in regulating NFκβ-P-p65. HUVEC treatment with TNF (24 hr) lead to an increase in P-p65 which was significantly decreased following genetic (siRNA of Panx1) or pharmacological (PxIL2P) inhibition of Panx1 (**Figure 4J-K**). Taken together, these data suggest that Panx1 channels in the plasma membrane act as a conduit for the movement of Ca^2+^ into the cytosol, leading to activation of NFκβ signaling, which acts as a feed-forward mechanism to promote the production and release of IL-1β.

## DISCUSSION

ATP release through Panx1 channels has been shown to be a strong signal for the recruitment of inflammatory cells in response to apoptosis (Chekeni et al., 2010) and activation of the NLR3P inflammasome (Albalawi et al., 2017; Lappas, 2014; Wu et al., 2017). However, the precise Panx1-mediated mechanisms controlling inflammation have never been fully elucidated. Here we demonstrate that EC-Panx1 is a direct target of the TNF signaling pathway, promoting plasma membrane localization of the channel, which facilitates the entry of extracellular Ca^2+^ into the cytosol. Importantly, we show that increases in intracellular Ca^2+^, resulting from enhanced Panx1 channel activity are directly associated with upregulated transcription and release of pro-inflammatory cytokines including IL-1β.

One of the primary findings in this study is the identification of a novel mechanism for the transcriptional regulation of Panx1. We observed that EC Panx1 is acutely sensitive to TNF stimulation, with maximal increases found in the low ng/mL range. Our data suggest that this is not a ubiquitous pathway, as increases in Panx1 expression were not found in TNF-treated SMCs or monocytes. Both SMCs and monocytes express TNF receptors and form inflammatory responses to TNF-stimulation (Gane et al., 2016; Jean-Charles et al., 2018; Warner and Libby, 1989). It is possible that cell-specific variances in gene or protein regulation in these cells may be due to differential regulation of receptor activation (Gane et al., 2016) or downstream pathways, e.g. SMC ubiquitin-specific protease 20 (USP20)-deubiquitinase activity which reduces NFκβ in SMC (Jean-Charles et al., 2018). There are few studies that have investigated the transcriptional control of Panx1 and there is currently limited data on pathophysiological mechanisms controlling Panx1 expression (Boyce et al., 2018). Jiang et al. reported increases in expression of Panx1 in mouse models of inducible stroke, which are associated with increased TNF-induced inflammation and tissue injury (Bokhari et al., 2014; Jiang et al., 2012; Liu and McCullough, 2011; Tuttolomondo et al., 2009; Tuttolomondo et al., 2014). In silico analysis has highlighted a number of transcriptional start sites within the rat Panx1 sequence, with binding sites for several transcription factors including CREB and ETV4 as well as factors downstream from IL-6 that have been identified (Dufresne and Cyr, 2014). Our RNA-seq data highlighted that TNF upregulates all NFκβ genes in HUVECs. TNF-induced Panx1 transcription was lost when HUVECs were pre-treated with inhibitors shown to block NFκβ-IKKB activity and TNF production (Gong et al., 2018; Kishore et al., 2003; Tobe et al., 2003). Our data strongly suggest that NFκβ pathways regulate EC Panx1 transcription.

We also found that increases in Panx1-transcription are followed by protein translation within 5 hours that was lost when cells were treated with the protein synthesis inhibitor CHX. The newly synthesized Panx1 proteins then translocate to the plasma membrane within 24 hours. Our surface biotinylation experiments highlight that the Panx1 isoform at the plasma membrane is primarily the complex-glycoprotein isoform (Panx1-Gly2). This is in keeping with previous studies suggesting that the Panx1-Gly2 is the predominant post-translationally modified form of Panx1 in the plasma membrane that forms hexameric membrane-channels (Boassa et al., 2007; Penuela et al., 2007). Pannexins are close family members of the connexin protein family which have a protein half-life of between 1-4 hours (Laird, 2006; Laird et al., 2017). An interesting observation in our study was that, once at the plasma membrane, the Panx1-Gly2 isoform is highly stable, unlike Cx43, and may persist for several days after the removal of TNF. This represents significantly longer protein stability than previously reported in experimental models by Boassa et al., and may highlight different protein recycling pathways between cell types (Boassa et al., 2007; Boassa et al., 2008). Despite having an extended residence time at the plasma membrane, our data also reveal that Panx1 channel activity may be dependent on continued stimulation by TNF for channel opening (as described in (DeLalio et al., 2019)), since [Ca^2+^]_i_ were reduced to baseline within 24 hours of removal of TNF. Thus, Panx1 functions in EC are regulated by TNF through multiple mechanisms including, synthesis, translation, trafficking and channel opening via phosphorylation.

Recent clinical trials have demonstrated that inflammation plays a key role in atherosclerotic disease development and that targeting specific cytokines may provide therapeutic benefits in high risk patients (Everett et al., 2013; Ridker, 2013, 2019; Ridker et al., 2019; Ridker et al., 2017; Ridker et al., 2012). In particular, the CANTOS trial demonstrated that the canakinumab, an anti-IL-1β therapeutic, reduces circulating levels of IL-1β in patients and decreases major adverse coronary events. Canakinumab treatments reduce IL-1β expression but also reduce expression of biomarkers including IL-6 and C-reactive protein (Ridker et al., 2017). However, targeting pathways associated with IL-6 expression using low-dose methotrexate, did not result in significant patient benefits (Chan and Cronstein, 2010; Cronstein et al., 1993; Ridker et al., 2019). This has led to the suggestion that atherosclerosis may be under the control of IL-1β and that targeting pathways controlling or controlled by IL-1β may have therapeutic benefit (Brown et al., 2015; Dinarello, 2010; Libby and Hansson, 2015; Ridker, 2013, 2019). Here we show that TNF treatment of HUVEC cells leads to release of cytokines including IL-1β, which was significantly reduced when Panx1 expression was knocked down via siRNA. Approaches using genetic (siRNA of Panx1) or pharmacological (PxIL2P) inhibition further identified that Panx1 expression and signaling can regulate IL-1β synthesis. These data suggest that Panx1 may be involved in multiple aspects of the synthesis and release of IL-1β in EC.

Our data provide further evidence that the expression of other inflammatory chemokines such as CXCL10 are controlled in a Panx1 specific manner. CXCL10 has been proposed to be an important inflammatory marker in atherosclerosis (Szentes et al., 2018; Zernecke et al., 2008), leading to the formation of vulnerable plaques in humans and mice (Heller et al., 2006; Segers et al., 2011). Strategies to lower CXCL10 expression can lead to reduced plaque formation and increased plaque stability (Segers et al., 2011). As previously described, it is possible that these increases correlate with IL-1β signaling pathways (Brown et al., 2015; Dinarello, 2010; Libby and Hansson, 2015; Ridker, 2013, 2019). However, the mechanisms through which Panx1 regulates these cytokines and chemokines remain to be fully elucidated.

The primary focus for signaling via Panx1 channels has been the release of ATP following channel opening (Chekeni et al., 2010; Lohman et al., 2015; Romanov et al., 2012), although other molecules are assumed to pass through these high conductance channels (Chiu et al., 2018; Vanden Abeele et al., 2006). Panx1 mediated ATP release has been associated with direct recruitment of inflammatory cells (Lohman et al., 2015), or through P2X receptor mediated pathways (Pelegrin, 2008). Receptor signaling via P2X receptors has leads to increases in the processing of pro-IL-1β to its mature form and with IL-1β release (Ainscough et al., 2015; Brough et al., 2003). Panx1-associated ATP release increases in Caspase 1 and Pro- IL-1β processing in human gestational tissues promoting (Lappas, 2014). In astrocytes, Panx1 mediated ATP release and signaling through the P2X7 channel, has been found promote activation of the NLRP3 inflammasome and the release of IL-1β (Albalawi et al., 2017). Our results show that TNF treatments (2.5-100 ng/mL) of HUVECs resulted in the release of approximately 10-20 nM ATP, which is similar to previous reports (Tozzi et al., 2019). Nonetheless, we found that adding exogenous ATP (data not shown) or the stable ATP isoform (ATP-γS) at these concentrations was not sufficient to stimulate IL-1β transcription. In previous studies, non-physiological levels of ATP (e.g., as high as 5 mM) induce IL-1β transcription (Cullen et al., 2015; Kanjanamekanant et al., 2013; Mehta et al., 2001). In keeping with these observations, we found that treatment of HUVECs with 100 μM ATP-γS did induce an increase IL-1β transcription, that was inhibited by the P2X inhibitor suramin. However, it should be noted that the ATP-induced IL-1β responses were significantly lower than those following TNF stimulation in our study. Furthermore, TNF-induced IL-1β and Panx1 expression in HUVECs was not altered by co-treatment with apyrase (to degrade ATP) or in the presence of suramin (to block P2X receptors). Taken together, these data suggest that ATP is not the primary mechanism for alterations in IL-1β synthesis in HUVECs following TNF treatment.

Panx1 channels are permeable to ions and molecules up to 1 KDa, and Vanden Abeele et al. previously demonstrated that Panx1 channels in the endoplasmic reticulum facilitates the movement of Ca^2+^ (Vanden Abeele et al., 2006). Here, we found that [Ca^2+^]_i_ was increased in HUVEC cells in response to TNF. Interestingly, this did not occur at earlier timepoints (5 hr) suggesting that kinase activation of TNF pathways is not involved. Rather, increases in intracellular Ca^2+^ were only found after long-term stimulation of up to 24 hr, a timepoint at which we demonstrate Panx1 channels are functional at the plasma membrane. To assess the source of increased intracellular Ca^2+^ we repeated the TNF stimulation experiment in Ca^2+^ free Krebs solution, which blocked the response, suggesting that increases in intracellular Ca^2+^ originated from outside the cell. While it is possible that other mechanisms serve to facilitate the entry of Ca^2+^ under these conditions, we provide several lines of evidence that suggest that Panx1 channels are directly permeable to Ca^2+^, these include no Ca^2+^ response to TNF when Panx1 is silenced by siRNA or the channel is inhibited using the Panx1 specific inhibitor peptide PxIL2P. Our data show that increases in intracellular Ca^2+^ are associated with IL-1β -synthesis, that can be ablated following reductions in intracellular Ca^2+^ using EGTA-AM and by blocking the Panx1 channel. Our data therefore suggest that plasma membrane Panx1 channels are permeable to, and facilitate increases in [Ca^2+^]_i_.

The NFκβ-p65 protein, is a downstream target of TNF signaling, and blocking its activation significantly alters TNF associated gene regulation, including IL-1β (Perrot-Applanat et al., 2011). Here, we have demonstrated that NFκβ activation plays a key role in early upregulation of Panx1 in EC, which promotes its membrane trafficking and channel opening. While Panx1 is a direct target of NFκβ activation, we further demonstrate that Panx1 signaling can enhance NFκβ activation. This is in keeping with studies by Wu et al that pointed to a role of Panx1 in the control of NFκβ activation (Wu et al., 2017). Our data highlight that TNF induces an increase in P-p65 activation, that is significantly reduced when Panx1 expression is knocked down by siRNA and when the channel is blocked in the presence of the PxIL2P peptide. In vitro, our results do not lead to a complete reduction in IL-1β expression, which may suggest that the role of Panx1-Ca^2+^ signaling is to amplify the NFκβ-mediated responses, through Ca^2+^-mediated phosphorylation of p65, as previously described (Martin et al., 2001). Thus, we propose that Panx1 facilitates a feed-forward signaling through Ca^2+^ leading to the transcriptional control of IL-1β.

Taken together our study highlights a novel reciprocal relationship between TNF and NFκβ signaling, which is regulated by Panx1 channel mediated control of intracellular Ca^2+^, leading to alterations in IL-1β synthesis and release by ECs.

## STAR ★ METHODS

Detailed methods are provided in the online version of this paper and include the following:

- **KEY RESOURCE TABLE**
- **CONTACT FOR REAGENT AND RESOURCE SHARING**
- **EXPERIMENTAL MODEL AND SUBJECT DETAILS**
  Primary cells and cell lines
- **METHOD DETAILS**
  Cell culture
  Cell transfection
  Cell treatment
  RNA extraction RT-qPCR and RNA seq
  Immunoblotting and membrane protein biotinylation
  Cytokine array
  Luciferase assay for ATP measurements
  Flow cytometric analysis of intracellular calcium and monocyte adhesion
- **QUANTIFICATION AND STATISTICAL ANALYSIS**

## METHODS

### KEY RESOURCES TABLE

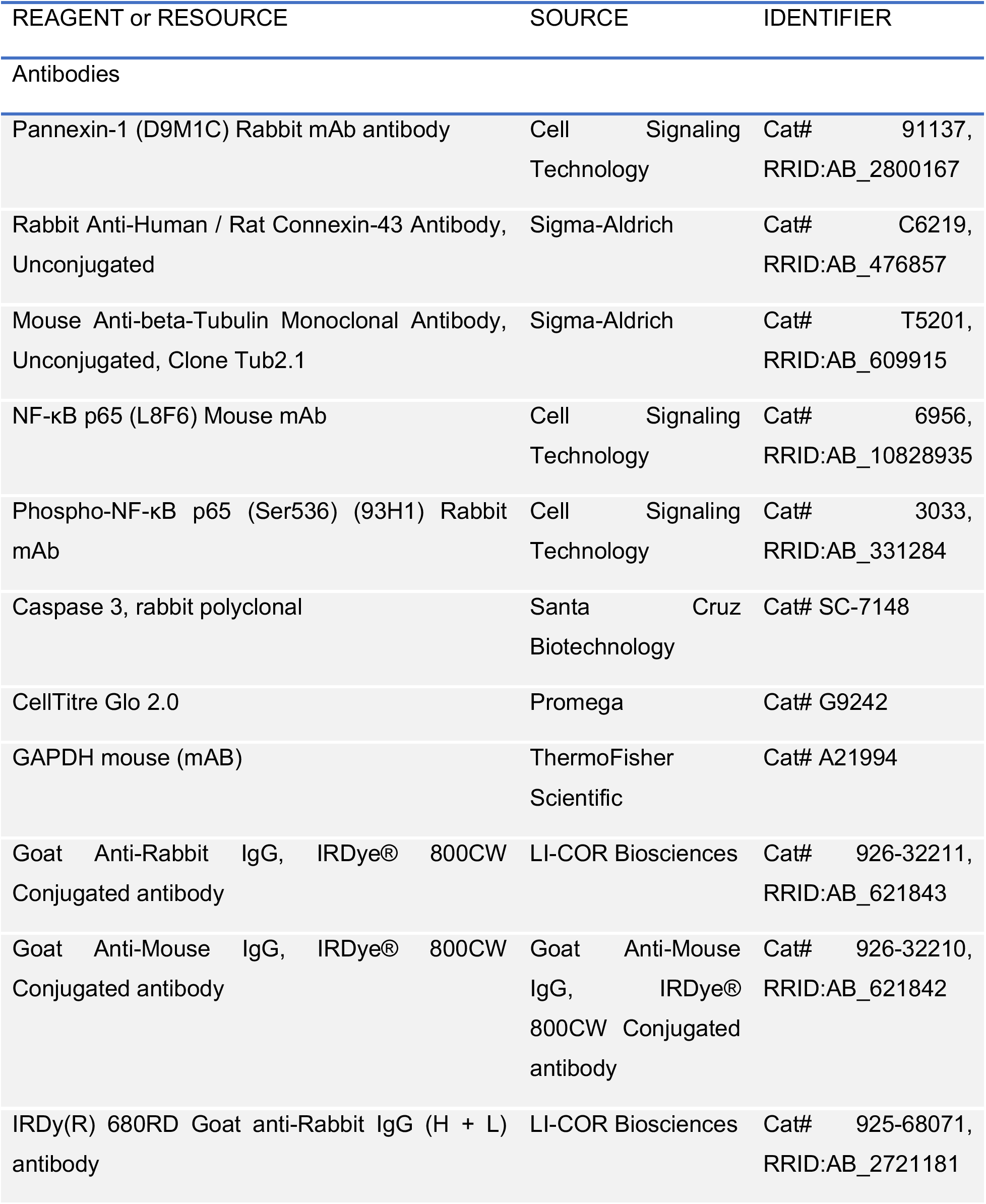
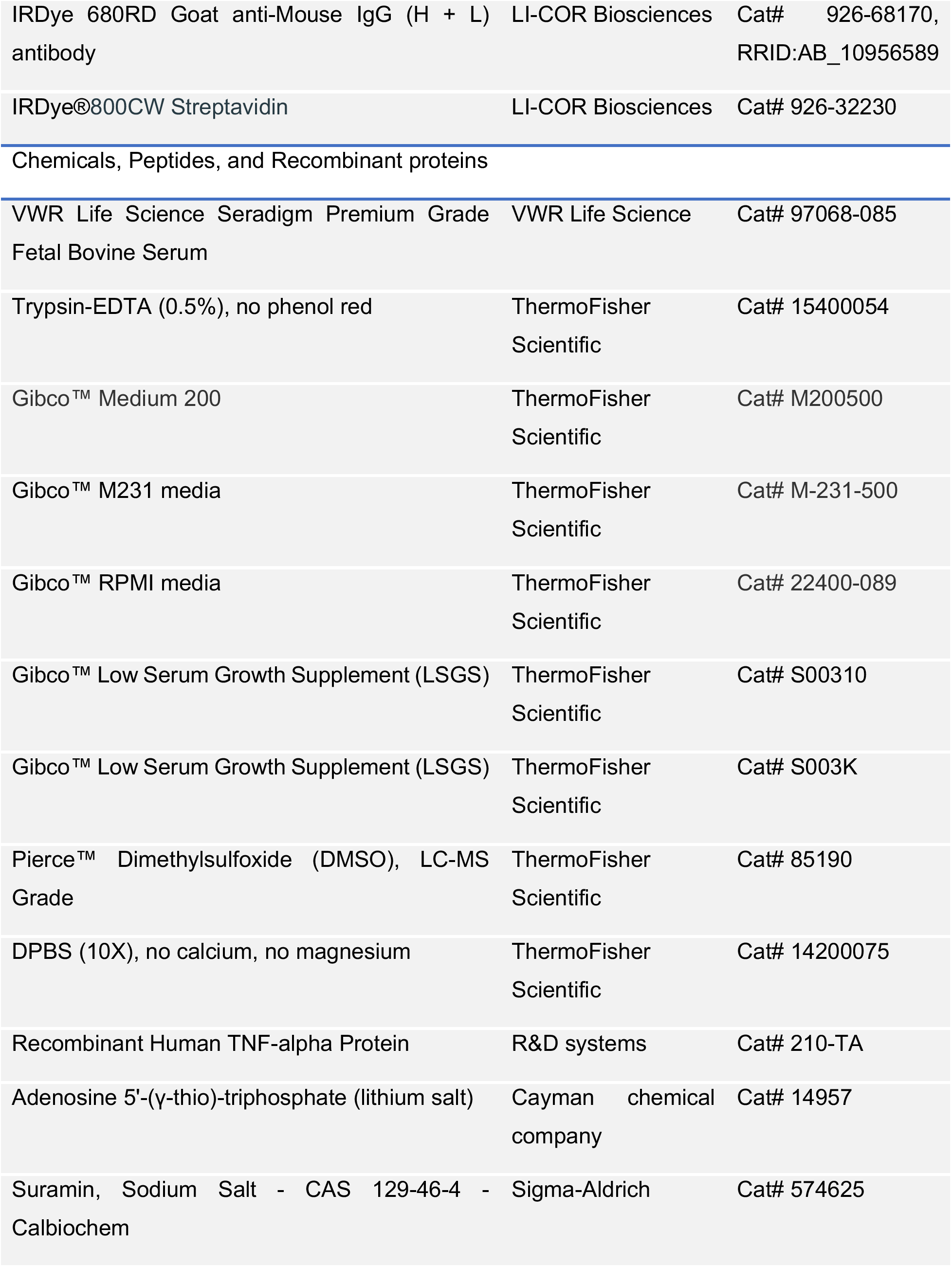
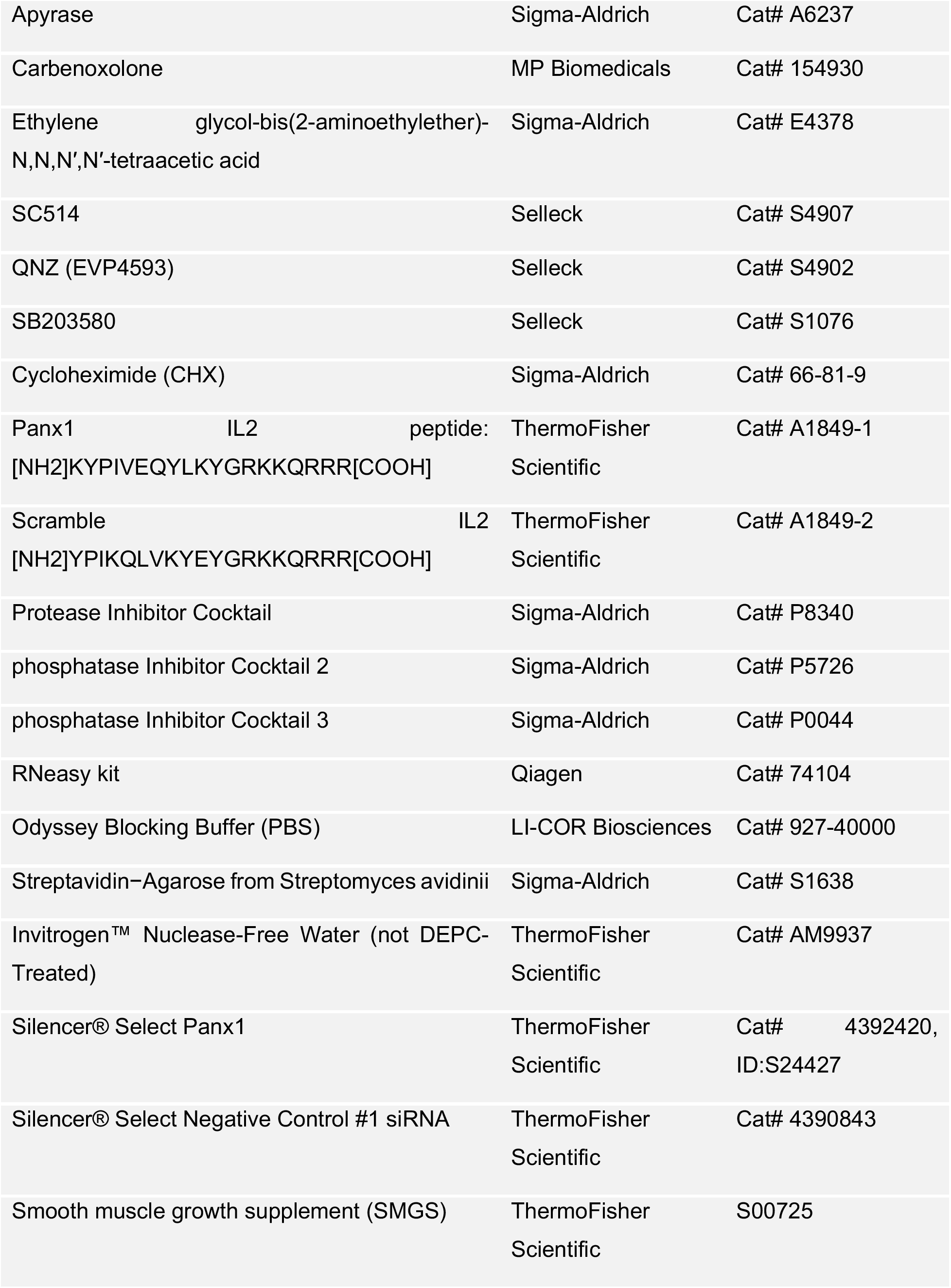
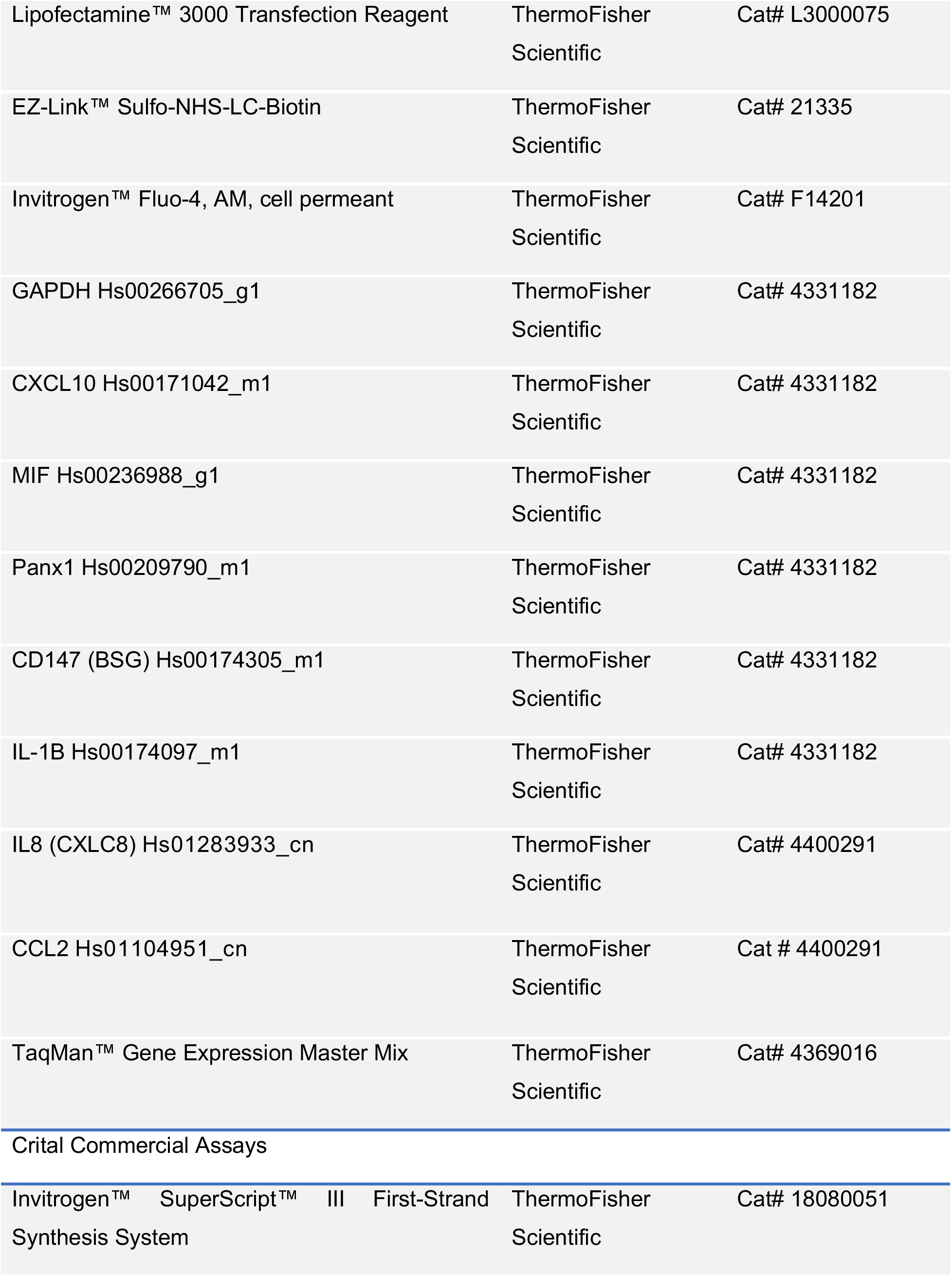
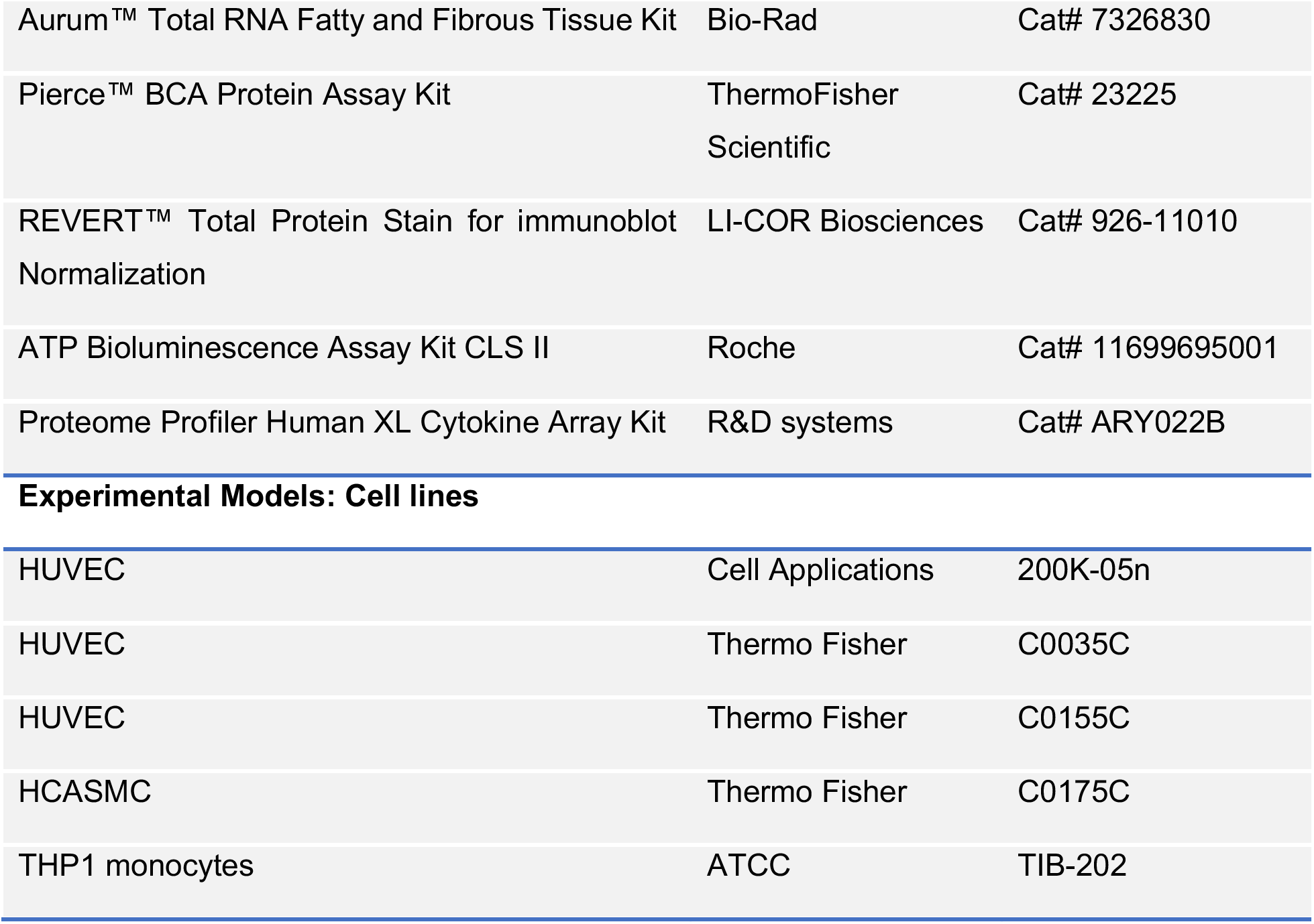

2 KEY RESOURCES TABLE
| REAGENT or RESOURCE | SOURCE | IDENTIFIER |
| --- | --- | --- |
| Antibodies |  |  |
| Pannexin-1 (D9M1C) Rabbit mAb antibody | Cell Signaling Technology | Cat# 91137, RRID:AB_2800167 |
| Rabbit Anti-Human / Rat Connexin-43 Antibody, Unconjugated | Sigma-Aldrich | Cat# C6219, RRID:AB_476857 |
| Mouse Anti-beta-Tubulin Monoclonal Antibody, Unconjugated, Clone Tub2.1 | Sigma-Aldrich | Cat# T5201, RRID:AB_609915 |
| NF-κB p65 (L8F6) Mouse mAb | Cell Signaling Technology | Cat# 6956, RRID:AB_10828935 |
| Phospho-NF-κB p65 (Ser536) (93H1) Rabbit mAb | Cell Signaling Technology | Cat# 3033, RRID:AB_331284 |
| Caspase 3, rabbit polyclonal | Santa Cruz Biotechnology | Cat# SC-7148 |
| CellTitre Glo 2.0 | Promega | Cat# G9242 |
| GAPDH mouse (mAB) | ThermoFisher Scientific | Cat# A21994 |
| Goat Anti-Rabbit IgG, IRDye® 800CW Conjugated antibody | LI-COR Biosciences | Cat# 926-32211, RRID:AB_621843 |
| Goat Anti-Mouse IgG, IRDye® 800CW Conjugated antibody | Goat Anti-Mouse IgG, IRDye® 800CW Conjugated antibody | Cat# 926-32210, RRID:AB_621842 |
| IRDy(R) 680RD Goat anti-Rabbit IgG (H + L) antibody | LI-COR Biosciences | Cat# 925-68071, RRID:AB_2721181 |
| IRDye 680RD Goat anti-Mouse IgG (H + L) antibody | LI-COR Biosciences | Cat# 926-68170, RRID:AB_10956589 |
| IRDye®800CW Streptavidin | LI-COR Biosciences | Cat# 926-32230 |
| Chemicals, Peptides, and Recombinant proteins |  |  |
| VWR Life Science Seradigm Premium Grade Fetal Bovine Serum | VWR Life Science | Cat# 97068-085 |
| Trypsin-EDTA (0.5%), no phenol red | ThermoFisher Scientific | Cat# 15400054 |
| Gibco™ Medium 200 | ThermoFisher Scientific | Cat# M200500 |
| Gibco™ M231 media | ThermoFisher Scientific | Cat# M-231-500 |
| Gibco™ RPMI media | ThermoFisher Scientific | Cat# 22400-089 |
| Gibco™ Low Serum Growth Supplement (LSGS) | ThermoFisher Scientific | Cat# S00310 |
| Gibco™ Low Serum Growth Supplement (LSGS) | ThermoFisher Scientific | Cat# S003K |
| Pierce™ Dimethylsulfoxide (DMSO), LC-MS Grade | ThermoFisher Scientific | Cat# 85190 |
| DPBS (10X), no calcium, no magnesium | ThermoFisher Scientific | Cat# 14200075 |
| Recombinant Human TNF-alpha Protein | R&D systems | Cat# 210-TA |
| Adenosine 5'-(γ-thio)-triphosphate (lithium salt) | Cayman chemical company | Cat# 14957 |
| Suramin, Sodium Salt - CAS 129-46-4 - Calbiochem | Sigma-Aldrich | Cat# 574625 |
| Apyrase | Sigma-Aldrich | Cat# A6237 |
| Carbenoxolone | MP Biomedicals | Cat# 154930 |
| Ethylene glycol-bis(2-aminoethylether)-N,N,N',N'-tetraacetic acid | Sigma-Aldrich | Cat# E4378 |
| SC514 | Selleck | Cat# S4907 |
| QNZ (EVP4593) | Selleck | Cat# S4902 |
| SB203580 | Selleck | Cat# S1076 |
| Cycloheximide (CHX) | Sigma-Aldrich | Cat# 66-81-9 |
| Panx1 IL2 peptide: [NH2]KYPIVEQYLKYGRKKQRRR[COOH] | ThermoFisher Scientific | Cat# A1849-1 |
| Scramble IL2 [NH2]YPIKQLVKYEYGRKKQRRR[COOH] | ThermoFisher Scientific | Cat# A1849-2 |
| Protease Inhibitor Cocktail | Sigma-Aldrich | Cat# P8340 |
| phosphatase Inhibitor Cocktail 2 | Sigma-Aldrich | Cat# P5726 |
| phosphatase Inhibitor Cocktail 3 | Sigma-Aldrich | Cat# P0044 |
| RNeasy kit | Qiagen | Cat# 74104 |
| Odyssey Blocking Buffer (PBS) | LI-COR Biosciences | Cat# 927-40000 |
| Streptavidin–Agarose from Streptomyces avidinii | Sigma-Aldrich | Cat# S1638 |
| Invitrogen™ Nuclease-Free Water (not DEPC-Treated) | ThermoFisher Scientific | Cat# AM9937 |
| Silencer® Select Panx1 | ThermoFisher Scientific | Cat# 4392420, ID:S24427 |
| Silencer® Select Negative Control #1 siRNA | ThermoFisher Scientific | Cat# 4390843 |
| Smooth muscle growth supplement (SMGS) | ThermoFisher Scientific | S00725 |
| Lipofectamine™ 3000 Transfection Reagent | ThermoFisher Scientific | Cat# L3000075 |
| EZ-Link™ Sulfo-NHS-LC-Biotin | ThermoFisher Scientific | Cat# 21335 |
| Invitrogen™ Fluo-4, AM, cell permeant | ThermoFisher Scientific | Cat# F14201 |
| GAPDH Hs00266705_g1 | ThermoFisher Scientific | Cat# 4331182 |
| CXCL10 Hs00171042_m1 | ThermoFisher Scientific | Cat# 4331182 |
| MIF Hs00236988_g1 | ThermoFisher Scientific | Cat# 4331182 |
| Panx1 Hs00209790_m1 | ThermoFisher Scientific | Cat# 4331182 |
| CD147 (BSG) Hs00174305_m1 | ThermoFisher Scientific | Cat# 4331182 |
| IL-1B Hs00174097_m1 | ThermoFisher Scientific | Cat# 4331182 |
| IL8 (CXLC8) Hs01283933_cn | ThermoFisher Scientific | Cat# 4400291 |
| CCL2 Hs01104951_cn | ThermoFisher Scientific | Cat # 4400291 |
| TaqMan™ Gene Expression Master Mix | ThermoFisher Scientific | Cat# 4369016 |
| Critical Commercial Assays |  |  |
| Invitrogen™ SuperScript™ III First-Strand Synthesis System | ThermoFisher Scientific | Cat# 18080051 |
| Aurum™ Total RNA Fatty and Fibrous Tissue Kit | Bio-Rad | Cat# 7326830 |
| Pierce™ BCA Protein Assay Kit | ThermoFisher Scientific | Cat# 23225 |
| REVERT™ Total Protein Stain for immunoblot Normalization | LI-COR Biosciences | Cat# 926-11010 |
| ATP Bioluminescence Assay Kit CLS II | Roche | Cat# 11699695001 |
| Proteome Profiler Human XL Cytokine Array Kit | R&D systems | Cat# ARY022B |
| Experimental Models: Cell lines |  |  |
| HUVEC | Cell Applications | 200K-05n |
| HUVEC | Thermo Fisher | C0035C |
| HUVEC | Thermo Fisher | C0155C |
| HCASMC | Thermo Fisher | C0175C |
| THP1 monocytes | ATCC | TIB-202 |

### CONTACT FOR REAGENT AND RESOURCE SHARING

Further information and requests for resources and reagents should be directed to and will be fulfilled by the Lead Contact, Scott R. Johnstone, PhD.

### EXPERIMENTAL MODEL AND SUBJECT DETAILS

#### Primary cells and cell lines

Human umbilical vein endothelial cells (HUVEC) and human coronary artery smooth muscle cells (SMC) were purchased from ThermoFisher and cell applications. Human THP1 cells were a kind gift Professor Zhen Yan, UVA.

### METHOD DETAILS

#### Cell Culture

HUVECs were grown in M200 media supplemented with the low serum growth kit and 20% FCS (20%-M200). For experiments involving TNF treatments, cells were incubated in M200 with low serum growth kit and 0.1% serum (0.1%-M200) for 24 hr. Human coronary artery smooth muscle cells (SMC, ThermoFisher) were grown in M231 media supplemented with the smooth muscle growth supplement (Thermo Fisher and 20% FCS. For experiments involving TNF treatments, cells were grown in M231 media containing 2% serum (2%-M231). Both HUVEC and SMCs were used within 16 population doublings to maintain primary phenotype. THP1 monocytes were grown in RPMI media supplemented with 10% FCS, 1% pen strep and 1% glutamine. All cells used are certified as mycoplasma free at start of experiments. All cell catalogue numbers are listed in the Key Resource Table.

#### Cell transfection

Endothelial cells were plated for expression (6 well plates) or ATP assays (24 well plates) until 70-80% confluent. Media was removed and replaced with 0.1%-M200 for 30 minutes prior to transfection. siRNAs targeting the human PANX1 gene (Life Technologies Panx1 silencer select 4392420-s24427; 50-GUUUGUGGGAG GUAUCUGAtt-30), Cx43 (Life Technologies GJA1 silencer select 4392420-s5758; 50-GGCUAAUUACAGUGCAGAAtt-30) or Control siRNAs (Life Technologies silencer select negative control 1, 4390843) or were transfected into ECs using Lipofectamine 3000. Media was changed after 24 hr and cells allowed to recover for 24 hr prior to treatment. siRNA knockdown for Panx1 was maximal at between 48-72 hr was confirmed by immunoblot and qRT-PCR.

#### Cell treatments

Prior to treatments, media was removed from cells and replaced with 0.1% -M200 for 24 hr. Media was then replaced with 0.1%-M200 containing 2.5 ng/mL TNF for up to 24 hr (as denoted per experiment). Where TNF dose responses were measured, the respective concentrations are denoted in the text. Inhibition of ATP (Apyrase, 1 UN/mL) and P2 activation (Surmain, 100μM) were performed by 30 minutes pre-incubation prior to addition of TNF for a further 24 hr. Inhibition of protein synthesis was performed by pre-incubation with 25 μg/mL cylohexamide.

##### Kinase inhibitors

Inhibitors of the IKK pathway SC514 (100 μM (Kishore et al., 2003)), and QNZ/EVP4593 (10 μM (Vigont et al., 2012)) and MAPK inhibitor SB203580 (10 μM (Adhikary et al., 2008)) were sourced from Selleck chemicals. All inhibitors were pre-incubated with cells for 3 hr prior to treatment with TNF for a further 24 hr.

#### RNA extraction RT-qPCR and RNA seq

Following treatments, media was removed, cells washed once in PBS, then 1mL of Trizol added prior harvesting by scraping. RNA was isolated using an RNA isolation kit (Aurum Total RNA isolation kit, #732-6830, BioRad) as per manufacturer protocol and cDNA synthesis performed using first strand synthesis kit (Thermo Fisher). Multiplex Taqman reactions were performed using 20ng of cDNA, Taqman primers (Thermo Fisher, see Key Resource Table) and Taqman gene expression mastermix (Thermo Fisher, #4369016). The internal control gene GAPDH was used for normalization and calculation of the delta CT values. All data are represented as delta-delta CT (2^-ΔΔCT) to define fold change from control values. For RNA-seq, Total RNA was isolated using an RNeasy kit (Qiagen) with an RNA free DNase step and quantity assessed on an Agilent 2100 Bioanalyzer. For experiments, three technical replicates (HUVEC no treatment and HUVEC TNF 2.5 ng/mL 24 hr) were sequenced with ribodepletion protocols. After sequencing 50 M reads were sampled from each replicate library and results analyzed by Glasgow Polyomics. Data from RNA-Seq is available through EMBL-EBI ArrayExpress, accession number E-MTAB-8299.

#### Immunoblotting and membrane protein biotinylation

Following treatments, all cells were harvested in cold lysis containing PBS (pH 7.4) containing: NaCl (125 mM); EDTA (5 mM); sodium deoxycholate (1%); triton X-100 (0.5%); sodium orthavanadate (500 μM); AEBSF (10 μM) and protease inhibitor cocktail (1:100, Sigma). All isolations were performed at 4 °C, samples were dounce homogenized 30 times on ice, incubated with rotation for 30 mins at 4 °C and centrifuged at 14,000g for 5 minutes. Cleared lysates were used for immunoblot analysis. Proteins samples were quantified by BCA assay prior to loading and equal loading confirmed using β-tubulin and total protein assays. Membranes were developed using Licor secondary antibodies anti-rabbit-700/800 and anti-mouse-700/800 (1:10,000) and imaged on a Licor Odyssey scanner. Expression analysis was performed using Image studio (Licor). Values were normalized to β-tubulin and changes calculated as a fold change compared to non-treated controls.

#### Cytokine array

Human cytokine array kit (R&D systems, proteome profiler XL ARY022) were used as per manufacturers protocols using 1mL of cleared culture media. Medias from 3 separate experiments under the same conditions were combined per reaction. Cytokine array membranes were developed using anti-streptavidin 800 on a Licor Odyssey scanner. Expression analysis was performed using Image studio (Licor). Values were normalized to the control spots on each blot and comparisons made to non-treated controls and expressed as a fold change from TNF control.

#### Luciferase assay for ATP release

ATP assays were performed as we have previously described (Lohman et al., 2015). Briefly, cultured EC were seeded in 24-well plates and grown to 70-80% confluency. Media was replaced with 0.1%-M200 media containing TNF (2.5 ng/mL) for 24 hr. On the day of the experiment cells were rinsed then incubated in 300 μL of fresh 0.1%-M200 media for 30 min at 37 °C, then incubated with the ectonucleotidase inhibitor ARL 67156 (300 μM, Tocris) for 30 min at 37 °C. Cells were then stimulated with recombinant human TNF (10 ng/mL) for 30 minutes. Following stimulation, 200 μL of the cell supernatants were collected, placed into pre-chilled tubes, centrifuged at 10,000 xg for 5 min and 50 μL of each sample was transferred to a white, opaque 96-well plate. ATP was measured using ATP bioluminescence assay reagents, CellTitre Glo 2.0 (Promega) or ATP Bioluminescence HSII kit (Roche). Using a FluoStar Omega luminometer, 50 μL of luciferin:luciferase reagent (ATP bioluminescence assay kit HSII; Roche) was injected into each well and luminescence was recorded following a 5-s orbital mix. For CellTitre Glo 2.0, reagent was mixed 50:50 with cleared HUVEC media. ATP concentration in each sample were calculated from an ATP standard curve. Data are presented either as calculated concentration or as a % change in ATP release from baseline (unstimulated cells) and expressed as mean±s.e.m. (N=5 independent experiments with triplicate measurements).

#### Flow cytometric analysis of intracellular calcium and monocyte adhesion

##### Monocyte adhesion assay

HUVECs were seeded on fibronectin coated plates and grown to 70% confluence. Cells were washed twice in warmed PBS and media changed for 0.1%-M200 for 24 hr. Media was changed for 0.1%-M200 with TNF (2.5ng/mL) for 24 hr. At the same time, human THP-1 monocytes were loaded with calcein-AM (0.1μM, Sigma) in RPMI for 30 mins. THP-1 cells were then centrifuged, and washed twice in PBS prior to being resuspended in 10% RPMI media for 24 hr. After 24 hr, TNF was removed from HUVEC by washing once in 0.1%-M200 and cells incubated in fresh 0.1%-M200 for 30 minutes. During this time, calcein loaded THP-1 cells were counted and resuspended to a concentration of 5×10^5^ cells/mL in 0.1%-M200. THP1 cells (100 μL, 5×10^4^) were added to each well for 4 hr at 37 °C. After 4 hr, the wells were washed gently two times with PBS to remove non-adherent cells. All remaining cells were then trypsinized and resuspended in 0.1%-M200 media and stored on ice for analysis by flow cytometry (BD FACScanto II). Gates for calcein stained THP1 and non-stained endothelial cells were defined and percentage of THP1 monocytes calculated from the total cells (endothelial cells and THP1 cells).

### QUANTIFICATION AND STATISTICAL ANALYSIS

1-way or 2-way ANOVA followed by Tukey or Dunnett post-test were used for comparisons between 3 groups and T-test used for comparisons of 2 treatment groups. A minimum of N=3 were used for all statistical analysis. In all analysis a P value of 0.05 is significant, * is P<0.05, *** is P<0.01, *** is P<0.001 **** is P<0.0001

## SUPPLEMENTAL INFORMATION

Supplemental information can be found online

## AUTHOR CONTRIBUTION

S.R.J., B.E.I. and M.K. conceived this study conceptualized and provided financial backing. S.R.J. and Y.Y. designed and executed the majority of the experiments. L.D., A.K.B., E.M., and J.M. performed extensive cell culture and immunoblotting. A.M., I.D. designed and performed RNA analysis for RNA-seq. M.M. performed RNA-seq and generation of ingenuity pathway data analysis. Y.Y. wrote the manuscript with input from X.H.S., M.K., B.E.I. and S.R.J.

## ACKNOWLEDGEMENTS

We thank Anita Impagliazzo for illustration. The UVA School of Medicine Flow Cytometry Facility was used for flow cytometric analysis. The University of Glasgow Polyomics core was used for RNA-seq and data interpretation and analysis. We thank Dr Graham Hamilton, Glasgow Polyomics for his input in experimental design and analysis of RNA-seq data.

## CONFILCT OF INTEREST

The authors have no conflicts to disclose.

## SUPPLEMENTAL FIGURE LEGENDS

**Supplemental Figure 1:**
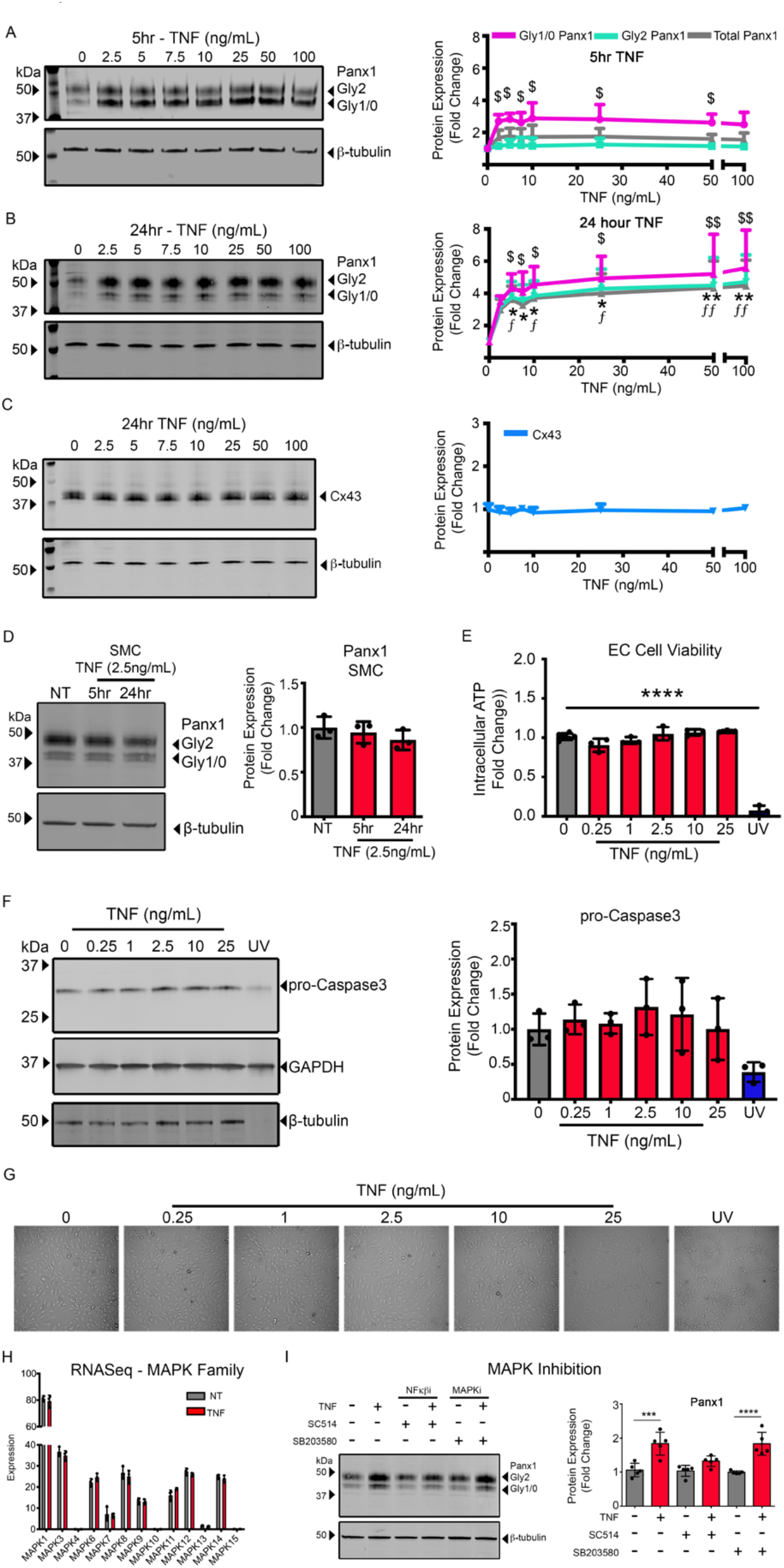
TNF increase of Panx1 in EC is not associated with activation of MAPK pathways or cell death. Protein lysates were isolated from HUVECs treated with a series of concentrations ranging from 2.5 ng/mL to 100 ng/mL TNF for either 5 or 24 hr. Non-treated cells were used as a control. Representative western blot analysis of Panx1 in HUVECs in response to doses of TNF for 5 hr (A) or 24 hr (B). In both A and B, Panx1 was analyzed as both bands (total Panx1) or as separate glycosylated species, Gly 2-Panx1 (upper band) or Gly 0/1-Panx1 (lower bands). Panx1 expression was normalized to β-tubulin and expressed as fold change from untreated (n=3). (C) Representative western blots of Connexin 43 (Cx43) in response to doses of TNF for 24 hr. Cx43 expression was normalized to β-tubulin and expressed as fold change from untreated (n=3). (D) Representative western blot of Panx1 in SMC treated with 2.5 ng/mL TNF for 5 and 24 hr. Panx1 expression was normalized to β-tubulin and expressed as fold change from untreated (n=3). (E) Intracellular ATP levels for HUVECs treated with doses of TNF (0.25, 1, 2.5, 10 and 25 ng/mL) for 24 hr or exposed under UV light for 10 minutes. Cell viability was expressed as fold change ATP luminescence in comparison to untreated cells (n=3). (F) Representative western blots for caspase 3 in HUVECs treated with a series of concentrations ranging from 2.5 ng/mL to 100 ng/mL TNF for 24 hr. Pro-caspase expression was normalized to β-tubulin and expressed as fold change from untreated cells (n=3). (G) Represent microscope images of HUVECs treated with a series of concentrations ranging from 2.5 ng/mL to 100 ng/mL TNF for 24 hr, show no evidence of cell death e.g. lower cell numbers or membrane blebbing. (H) RNA-Seq results of HUVECs treated with TNF (2.5 ng/mL). The expression of thirteen genes in MAPK family are shown with each bar represents mean±SD for triplicates (n=3). (I) Representative western blot of TNF (2.5 ng/mL)-induced HUVECs in presence or absence of inhibitors: inhibitor of nuclear factor kappa-B kinase-2 (IKK2) 100 μM SC514 (n=5) or MAPK inhibitor 10 μM SB203580 (n=3). In A-E, Statistical analyses were performed by one-way ANOVA with Dunnett’s’ multiple comparison test, * indicates comparison to not treated Total-Panx1, $ indicates comparison to not treated Gly1/0-Panx1, F indicates comparison to not treated Gly2-Panx1. ^*/$/f^P<0.05, ^$$/ff^P<0.01, ****P<0.001. In I, Statistical analyses were performed by two-way ANOVA with Tukey’s multiple comparison test, ***P<0.001, ****P<0.001.

**Supplemental Figure 2:**
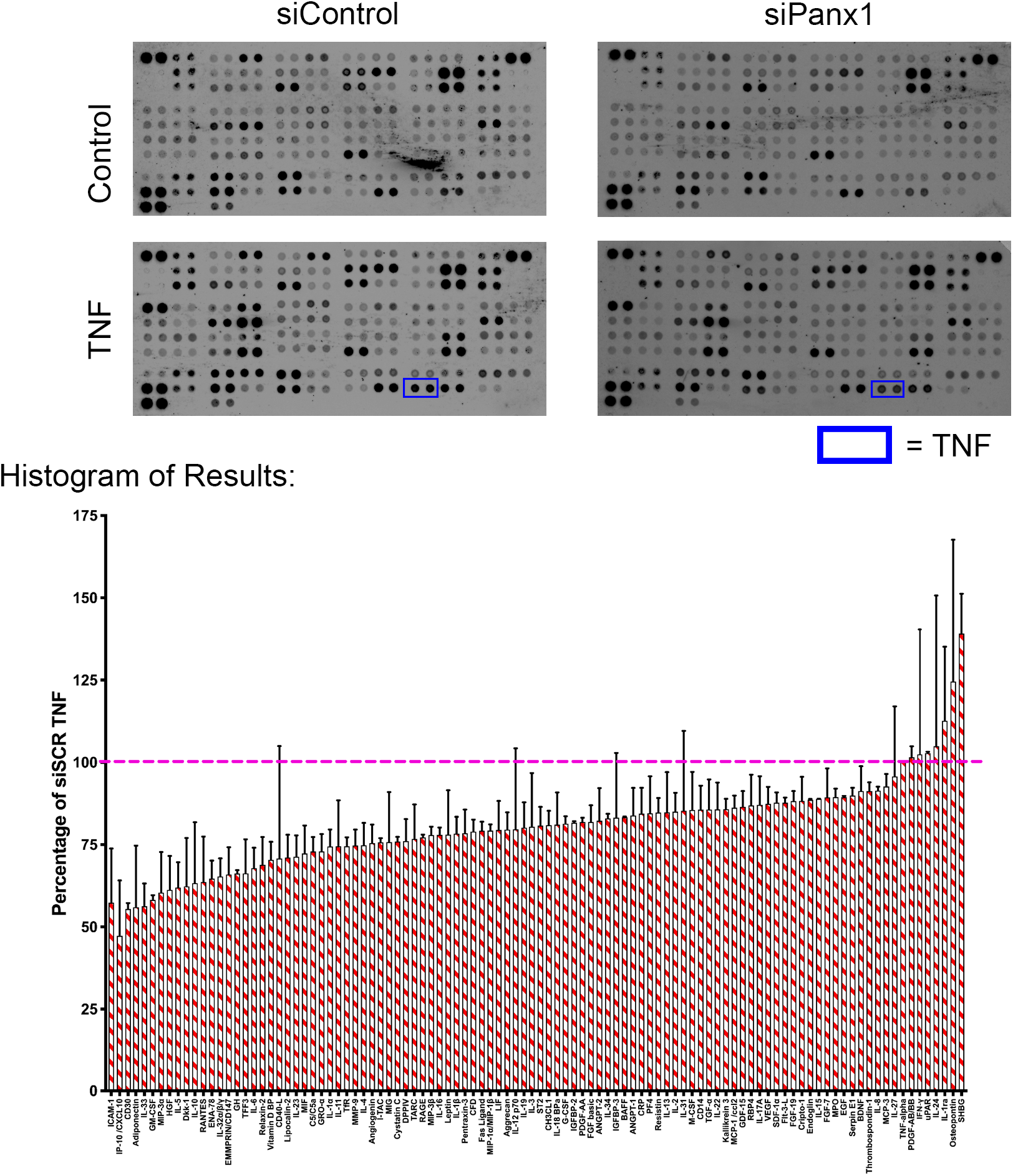
Cytokine array of HUVEC treated with TNF. Representative human cytokine array membranes (siControl and siPanx1 with or without TNF) show 102 biomarkers spotted in duplicate and arranged in a grid format. HUVECs transfected with control siRNA (siControl) or Panx1 (siPanx1) for 48 hr followed with TNF (2.5 ng/mL) treatment for 24 hr. Media from 3 technical replicates were pooled and incubated with array membranes. Images were visualized using LICOR Odyssey imaging system. Differences between TNF-treated siControl and siPanx1 were measured by LICOR in 2 independent experiments. The fluorescence intensity of cytokines were normalized to reference values on blots and expressed as the percentage of siControl TNF in the histogram (n=2).

**Supplemental Figure 3:**
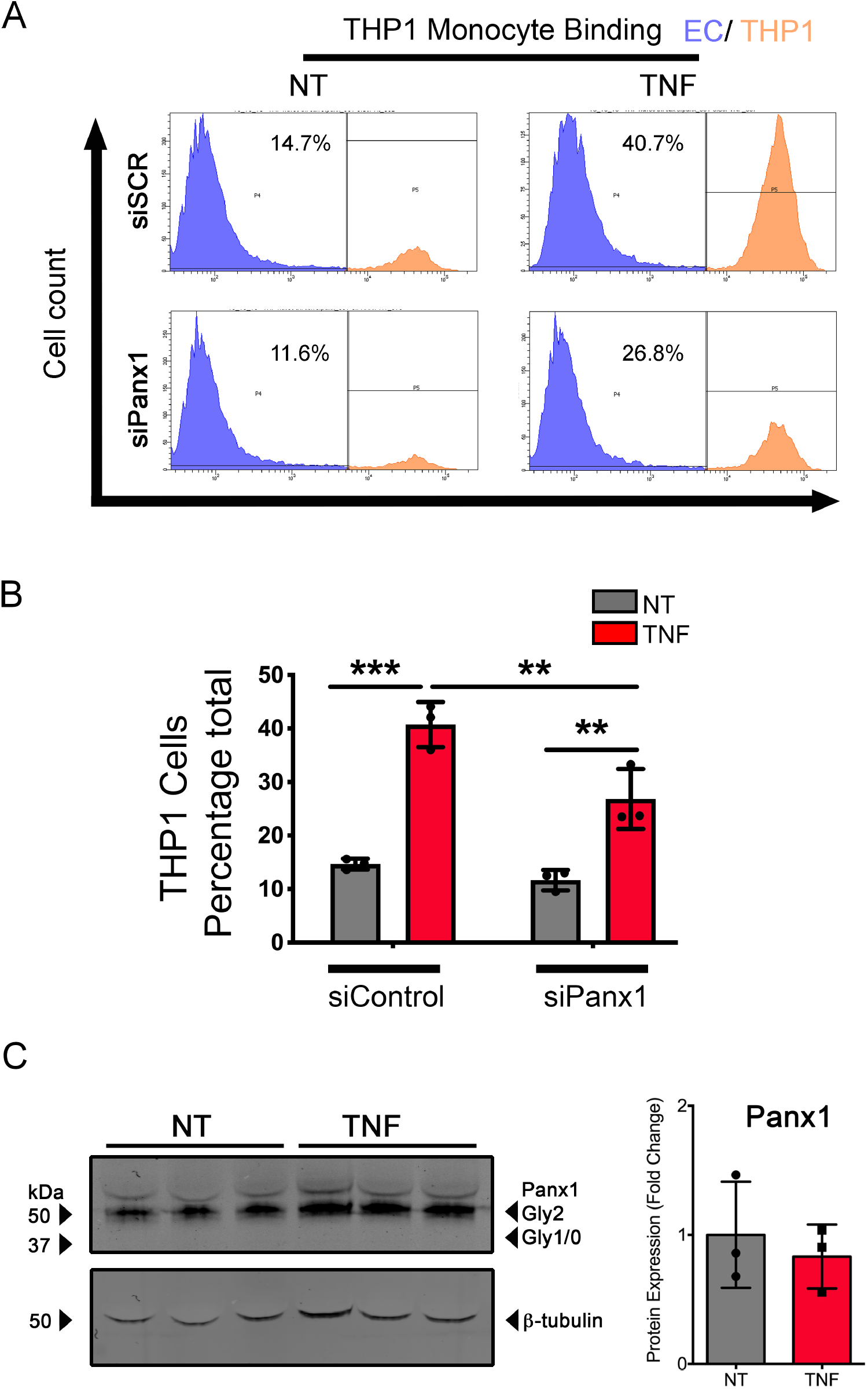
Panx1 regulates TNF associated leukocyte binding. Representative histograms from flow cytometry of calcein loaded leukocyte binding with HUVEC transfected with either control siRNA (siControl) or Panx1 siRNA (siPanx1) for 48 hours prior to TNF (2.5 ng/mL) stimulation for 24 hr. (A) Representative histograms highlighting percentage calcein-THP1 cells bound to HUVECs in each group. (B) Percentage of leukocyte binding in triplicates. Statistical analyses were performed by two-way ANOVA with Tukey’s multiple comparison test, *P<0.05, **P<0.01, ***P<0.001, ****P<0.001. (B) Western blot analysis of Panx1 in human monocytes (THP1) treated with 2.5ng/mL TNF for 24 hr. Total protein expression values of Panx1 were normalized to β-tubulin and expressed as fold change (n=3). Statistical analyses were performed by Student’s test.

